# Capturing the Mutational Dynamics of SARS-CoV-2 with Graphs

**DOI:** 10.1101/2025.09.02.673804

**Authors:** Badhan Das, Lenwood S. Heath

## Abstract

The rapid evolution of SARS-CoV-2 presents significant challenges for modeling viral dynamics, driven by lineage diversification and region-specific mutation patterns. While phylogenetic trees are traditionally used for evolutionary inference, the massive volume of SARS-CoV-2 genomic data, with many similar sequences and few distinguishing mutations, poses computational and methodological limitations. The quasispecies theory instead models viral evolution as a cloud of mutants, motivating a graph-based representation that better captures the complexity of mutational events. Geographic variation adds another critical layer to this complexity. Mutation trends often differ across regions due to local transmission dynamics, host population structures, and selective pressures.

In this study, we present the Mutation Learning Graph (MLG), a directed graph framework that organizes SARS-CoV-2 variants based on their cumulative mutation profiles relative to the reference genome (NC_045512.2), thereby capturing the dynamics of mutation propagation. This structure captures fine-grained mutational transitions and encodes plausible evolutionary relationships among variants. To construct these graphs, we introduce an alignment-aware mutation profiling method and a novel Ancestor-Joining algorithm, which incorporates ancestral variants as inferred intermediate nodes to connect observed genomes through biologically coherent mutational paths. We generate MLG datasets for ten geographically and epidemiologically diverse regions and benchmark them on two graph-based tasks: node-level lineage classification and edge-level mutational transition prediction. Using baseline graph neural network architectures (GCN, GraphSAGE, GAT, GGNN, VGAE), we demonstrate how mutation-centric graph structures expose key biological challenges, such as lineage imbalance and location-specific mutation spectra. For node classification, GraphSAGE and GGNN consistently achieved high accuracy (up to 0.96) and AUROC (up to 0.98). In contrast, VGAE and GraphSAGE led the way in link prediction, with AUPRCs of up to 0.96. These results highlight the effectiveness of MLG for capturing biologically meaningful mutation patterns and underscore the importance of localized, mutation-aware modeling for predicting viral mutations and future variant emergence.

## 1 Introduction

The emergence of severe acute respiratory syndrome coronavirus 2 (SARS-CoV-2) in late 2019 triggered a global pandemic with profound public health, social, and economic consequences. As the causative agent of coronavirus disease 2019 (COVID-19), SARS-CoV-2 spread rapidly across the globe, resulting in more than 770 million confirmed cases and nearly 7 million deaths as of early 2024 [53]. A significant challenge in managing the pandemic has been the continuous evolution of the virus, driven by the high mutation rate typical of RNA viruses [10]. SARS-CoV-2, as with other coronaviruses, possesses an RNA genome that mutates frequently during replication, giving rise to new genetic variants with altered transmissibility, immune escape properties, or virulence [20, 29]. Monitoring and analyzing these mutations is crucial for understanding viral evolution, assessing the effectiveness of vaccines and therapeutics, and informing public health responses.

Although phylogenetic trees have been the standard for reconstructing viral evolution, they impose strict bifurcating structures that fail to account for key biological complexities in viral evolution, such as recurrent mutations, back mutations, homoplasies, and convergent evolution [3, 7, 13, 14, 24, 36, 46, 47, 56, 57]. The constantly growing SARS-CoV-2 genomic data pose a significant computational challenge for phylogenetic analysis [21, 34]. According to Morel et al. [36], inferring a reliable phylogeny from GISAID [43] data is hard due to the vast quantity of sequences and the limited number of mutations.

Viral evolution is shaped by replication errors, selection pressures, and transmission dynamics [23, 39]. These processes generate a complex, heterogeneous cloud of viral genomes known as a viral quasispecies [1, 9], wherein variants coexist and compete within a single host. Quasispecies theory outlines the development of an infinite population of asexual replicators with high mutation rates [11]. This concept, grounded in molecular evolution theory [11, 12], highlights the interplay between genetic diversity and fitness landscapes. Many variants are lost due to transmission bottlenecks [5], while others fix due to selective advantages, such as the D614G spike mutation [50]. By analyzing shared and unique mutations, researchers can reconstruct plausible evolutionary relationships and uncover lineage dynamics, including competition and replacement [8, 33].

Recent efforts have employed machine learning to analyze the trends and effects of SARS-CoV-2 mutations. Hossain et al. [22] used LSTMs to predict mutation frequencies, while Li et al. [30] applied CNNs to classify variants based on dinucleotide composition. Serna GarcÍa et al. [41] used GPT-2 to extract literature-based mutation-effect associations. Maher et al. [32] forecasted the short-term spread of mutations using protein language models. In contrast, Taft et al. [44] and Han et al. [19] developed deep learning models to predict ACE2 binding and immune escape. Other works have emphasized structural modeling [2, 28, 37, 49, 52]. While these approaches capture aspects of mutation impact or prevalence, they typically do not model how mutations propagate over time or how variants can be related through directed mutational paths, limiting their use for forecasting novel variant emergence.

Understanding viral evolution requires examining mutations at a fine-grained level. In this study, we observe the granular details of SARS-CoV-2 mutations to identify patterns that characterize the emergence and propagation of variants. Such detailed mutation-level analysis is essential for capturing the evolutionary dynamics of the virus, particularly in the face of complex phenomena such as parallel evolution, back mutations, and regional adaptation. This fine-scale profiling is critical for portraying how variants emerge, accumulate mutations, and evolve. Geographic location also plays a key role, as mutation patterns vary across regions due to differences in transmission dynamics, host genetics, selective pressures, and public health interventions. Therefore, building predictive models for future mutations requires not only mutation-aware learning but also a location-specific perspective that captures regional evolutionary trends.

To capture the viral complexities and mutational dynamics, we introduce the Mutation Learning Graph (MLG). This directed graph organizes viral mutants as vertices based on their cumulative mutation profiles relative to the reference genome (NC_045512.2) [54], capturing the propagation of mutations as edges among variants over time. The resulting graph structures naturally enable the use of graph-based machine learning models for predictive analysis, such as classifying variant lineages or forecasting mutational transitions. To build this graph, we propose an alignment-aware mutation profiling approach and a novel Ancestor-Joining algorithm that infers unsampled intermediates or ancestral variants connecting observed variants through plausible mutational transitions, thereby preserving biological directionality. Our framework also accounts for back mutations [4, 42, 46], a less frequent yet biologically plausible event. We have curated ten region-specific SARS-CoV-2 data sets from Egypt, Iran, Nigeria, Bangladesh, Queensland (Australia), China, Estonia, Wyoming (USA), Chile, and South Africa, spanning diverse geographic, epidemiological, and evolutionary contexts, and demonstrating how these data sets encode meaningful patterns of mutation propagation. These graphs feature rich node and edge attributes, making them immediately compatible with standard graph-based ML models and enabling downstream analysis through graph machine learning.

To validate the utility of our proposed graph, we benchmark a suite of graph-based baseline models, including GCN, GraphSAGE, GAT, GGNN, and VGAE, on two key prediction tasks: lineage classification (node-level) and mutational edge prediction (link-level), demonstrating the practical challenges of heterogeneous and imbalanced distributions of lineages, regional diversity, and bidirectional mutational behavior. We also provide code and detailed documentation to facilitate future research and data set generation for SARS-CoV-2 or other viruses. While our method may not capture the exact history of evolution, it offers a robust mutation-aware structure that integrates genomic, temporal, and topological signals to support viral evolutionary modeling. This generalizable framework can be readily extended to other viral outbreaks, including pandemics or epidemics, providing a scalable foundation for designing curated mutation-centric datasets to train graph-based models.

## 2 Methods

### 2.1 Data Preprocessing

We obtained the SARS-CoV-2 reference genome (NC_045512.2) from NCBI [54] and downloaded genome sequences from GISAID [43] of ten geographic locations: Egypt, Iran, Nigeria, Bangladesh, Queensland (Australia), China, Estonia, Wyoming (USA), Chile, and South Africa. Only sequences labeled as *complete, high-coverage*, and with valid *collection dates* were retained to ensure data quality. To support accurate mutation modeling, we excluded any sequences containing ambiguous nucleotides (`N`) in the coding region.

For each location, the set of *m* high-quality genome sequences, along with the reference genome, is aligned using MAFFT (Multiple Alignment using Fast Fourier Transform) [25]. To reduce noise, we removed the untranslated regions (UTRs), positions [1 *−* 265] (5^*′*^ UTR) and [29675 − 29903] (3^*′*^ UTR), focusing instead on the coding region [266 −29674], which spans all annotated open reading frames (ORFs), including ORF1ab, S, E, M, N, and accessory genes.

To eliminate redundancy due to repeated sampling of the same variant, we deduplicate sequences by retaining only the earliest sample with identical alignments, based on the collection date. This process reduces the data set to *k* unique genome alignments, where *k* ≤ *m*, thereby preserving evolutionary diversity while minimizing duplication. A schematic overview of the preprocessing workflow is presented in Fig. 1.

**Figure 1:**
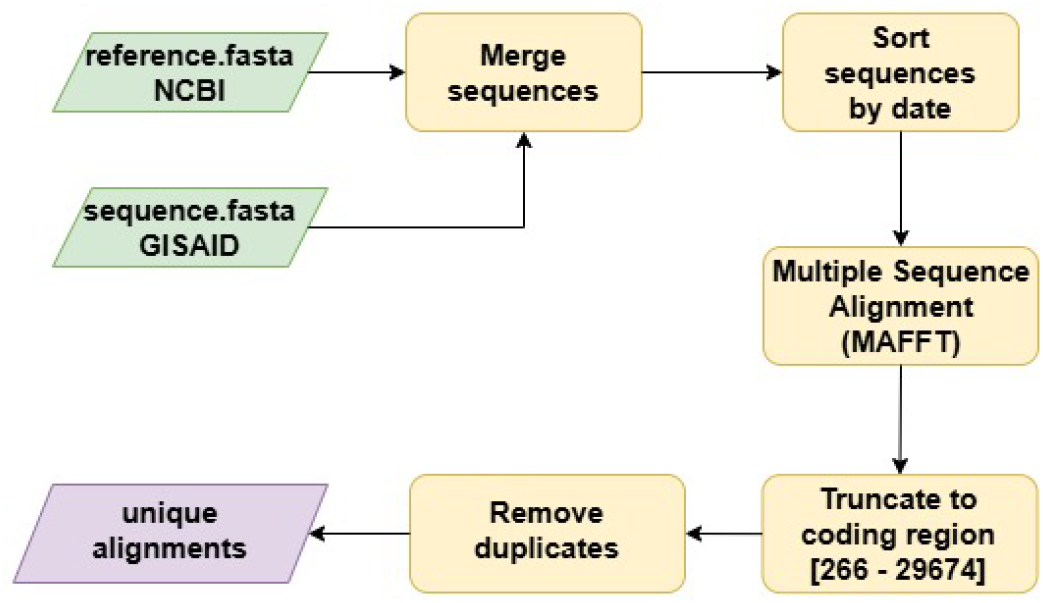
Data preprocessing pipeline

### 2.2 Mutation Function

Multiple sequence alignment enables direct, position-wise comparison of genomes by aligning sequences to a reference and standardizing lengths with gap insertions. This ensures that each alignment position maps to the same genomic coordinate, simplifying and organizing mutation detection. Substitutions appear as mismatches, and insertions and deletions (indels) appear as gaps (-).

Let Σ = {*A, C, G, T*, −} denote the nucleotide alphabet including gaps. Let the set of unique alignments be *S* = {*s*^1^, *s*^2^, …, *s*^*k*^}, where each aligned genome sequence *s*^*i*^ ∈ Σ^*n*^ is of length *n*, and the aligned reference genome is denoted by

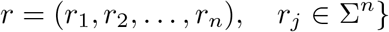

Each variant *s* = (*s*_1_, *s*_2_, …, *s*_*n*_) *∈* Σ^*n*^ represents a genome sequence aligned to the reference. We define the mutation function ℳ (*s*) as,

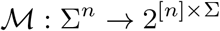

that returns the mutation set for a given variant sequence *s* with respect to the reference genome *r* as

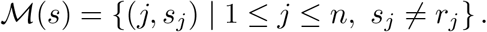

That is, ℳ (*s*) contains all position-nucleotide pairs (*j, s*_*j*_) where the nucleotide in *s* differs from the aligned reference *r*. These differences include substitutions and gap-induced indels, and form the foundation for mutation-based analyses.

### 2.3 Edit-distance Matrix

Levenshtein distance [51] or edit distance measures the minimum number of edit operations (insertions, deletions, or substitutions) required to transform one string into another. For aligned sequences of equal length, this reduces to the Hamming distance [18], which enables direct, position-wise comparison of nucleotide differences.

Let ℳ (*a*) and ℳ (*b*) denote the mutation functions of two sequences *a* and *b*, where each maps to a set of mutated positions and nucleotide pairs. Let,

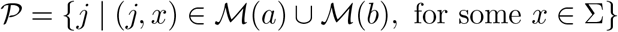

as the set of all genomic positions having a mutation in either *a* or *b*. Then, the mutation-aware edit distance between *a* and *b* is defined as:

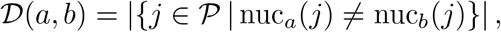

where

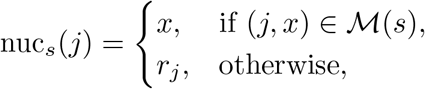

where *s* ∈ *S*, and *r*_*j*_ is the nucleotide at position *j* in the aligned reference genome, *r*.

We first compute the mutation function ℳ (*s*) for each sequence *s* ∈ *S*, where each ℳ (*s*) ⊆ [*n*] × Σ records positions where *s* differs from the reference genome *r*. This step requires 𝒪 (*nk*) time. Pairwise distances are then computed for the edit-distance matrix, *E*, of size *k × k*, using the symmetric difference ℳ (*a*) Δ ℳ (*b*), with time complexity (𝒪|ℳ (*a*) | + |ℳ (*b*) |) per pair. Letting 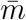 denote the average number of mutations per sequence (typically 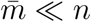), the total cost becomes 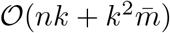, offering substantial scalability for large viral genome data sets like SARS-CoV-2.

We compare three strategies: classical Levenshtein (edit) distance, Hamming distance, and our proposed mutation-aware edit distance, for computing pairwise distances and edit-distance matrix for *k* aligned genome sequences of length *n*. Classical Levenshtein distance requires 𝒪 (*n*^2^) time per pair, resulting in a total complexity of 𝒪 (*k*^2^*n*^2^). Hamming distance improves this to *O*(*n*) per pair and 𝒪 (*k*^2^*n*) in total for the complete matrix. A summary of these complexities is provided in Table 1.

**Table 1:** Time complexities for computing pairwise distances and edit-distance matrix among *k* aligned sequences of length *n*. Here, *m* and *m*^*′*^ are mutation counts in the compared sequences, and 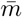 is the average number of mutations, where 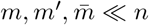.

| Method | Single Pair | Full Matrix |
| --- | --- | --- |
| Edit Distance (Levenshtein) | $\mathcal{O}(n^2)$ | $\mathcal{O}(k^2n^2)$ |
| Hamming Distance (Aligned) | $\mathcal{O}(n)$ | $\mathcal{O}(k^2n)$ |
| Mutation Set-based (Proposed) | $\mathcal{O}(m + m')$ | $\mathcal{O}(nk + k^2\bar{m})$ |

### 2.4 Mutation-Similarity Matrix

For each aligned sequence *s* ∈ *S*, we computed its mutation function ℳ (*s*), which returns the set of genomic positions and nucleotide pairs where *s* differs from the reference genome. Using these sets, we construct a mutation-similarity matrix, Π, of size *k* × *k*, where each entry (*i, j*) represents the number of shared mutations between sequences *s*^*i*^ and *s*^*j*^, computed as |ℳ(*s*^*i*^) ∩ ℳ(*s*^*j*^)|. Since each sequence is trivially most similar to itself, the diagonal entries naturally contain the highest values. However, to ensure that the similarity computation emphasizes relationships between distinct variants rather than self-comparisons, we set all diagonal entries to zero. This matrix captures pairwise mutational overlap across the aligned genomes and serves as a critical input for constructing Mutation Learning Graphs (MLGs) and for enabling mutation-informed downstream analyses.

### 2.5 Definition of the Mutation Learning Graph (MLG)

We define the Mutation Learning Graph (MLG) as a directed graph *G* = (*V, E*) in which:

- Each node *v* ∈ *V* represents a unique viral genome sequence, either *observed* in the dataset or *inferred* as a plausible ancestral or intermediate variant.
- Each directed edge (*u, v*) ∈ *E* represents the mutational transition from genome *u* to genome *v*, where the mutation set *M* [*u*] is a subset of *M* [*v*] relative to the reference genome NC_045512.2.
- Edges may be annotated as *back mutations* when the collection date of *v* precedes that of *u* despite *M* [*u*] ⊂ *M* [*v*].
- Node and edge attributes include: mutation sets, collection dates, lineage labels (observed or predicted), edit distances, and mutation similarities.

### 2.6 Mutation Learning Graph

We propose a two-phase Ancestor-Joining algorithm (Algorithm 1) to construct a Mutation Learning Graph. This directed graph highlights the mutational connectivity between genome variants. This algorithm maintains mutational reachability by inserting *inferred* nodes to connect observed nodes when necessary, producing a connected, cohesive, biologically plausible graph. Each node is either an *observed* variant or an *inferred* ancestor; each edge encodes the precise nucleotide changes required to reach one variant from another. The entire pipeline for building MLG is illustrated in Fig. 2. The pseudocode in Algorithm 1 presents a high-level view of the construction process, with several helper functions (e.g., `kNN_Group, Bridge_Components`, and `Expand`) implemented with additional logic for biological validity and graph completeness. Detailed descriptions of these helper functions and implementation-specific choices are provided in the Supplementary Material.

**Figure 2:**
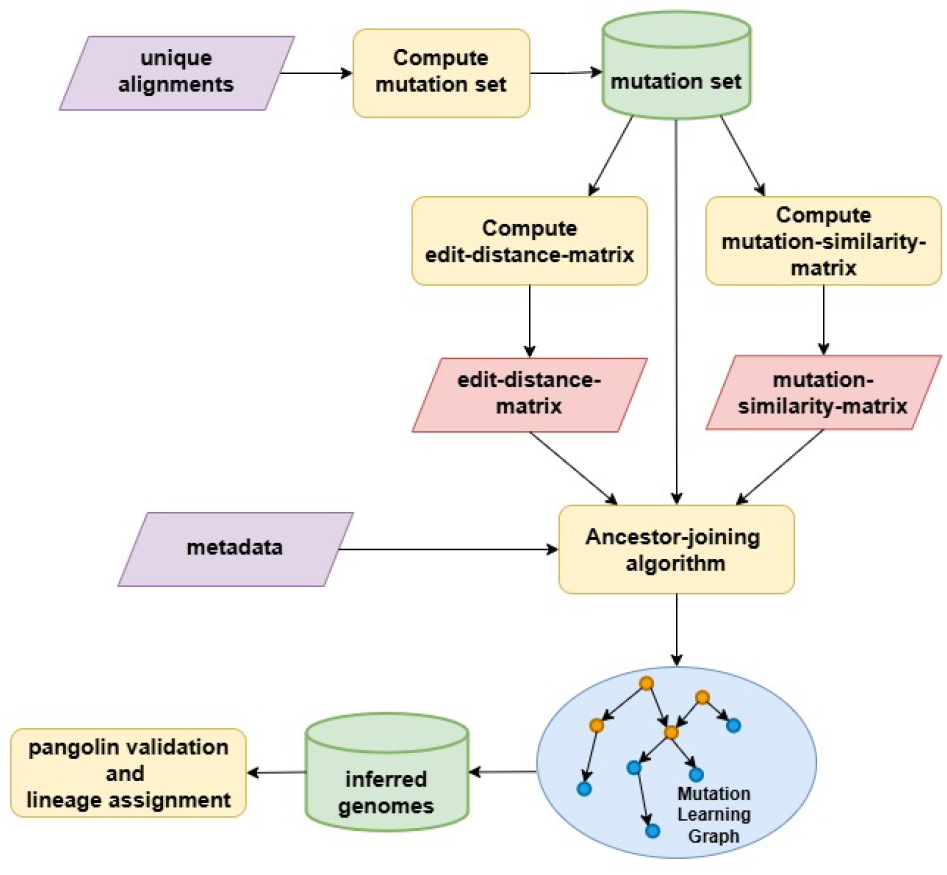
Pipeline for building MLG using the Ancestor-Joining algorithm.

#### Phase A (Local *k*-NN Pass)

For every variant, we retain its top *k* = 5 highest similarity neighbors (i.e., row-wise *k*-NN on the similarity matrix Π). Inside this *k*-NN mask, we select the cell(s) that jointly minimize the edit distance *E* and maximize the similarity Π.

Formally, we minimize the lexicographic pair (*E*[*i, j*], −Π[*i, j*]), prioritizing neighbors with the smallest edit distance and, among ties, the highest similarity. This strategy accounts for the frequent occurrence of dense clusters of genomes that differ at only a few nucleotide positions, typical of SARS-CoV-2 mutation profiles.

All neighbors tied under this criterion are grouped into a single equivalence class *g* with the current variant. Let 𝒞 be the intersection of their mutation sets. If 𝒞 ≠ ∅ and a node (observed or inferred) already carries 𝒞, that node becomes the parent of every variant in *g*. Otherwise, the algorithm introduces an inferred ancestor with mutation set 𝒞 and links it to each member of *g*. This step encodes strong local ancestry signals while preserving mutational coherence.

#### Phase B (Component Bridging)

Following Phase A, the graph may consist of multiple weakly connected components. To ensure overall connectivity, we identify inter-component connections using a global max-heap [6] containing all remaining candidate edges ranked by similarity Π, and maintain component membership with a disjoint-set (union–find) data structure [45].

At each iteration, we extract the highest-similarity pair (*u, v*) such that *u* and *v* belong to different components. These nodes are then connected using the same logic as in Phase A: if their mutation sets have a non-empty intersection 𝒞, and no node already carries 𝒞, we introduce a new inferred ancestor with mutation set 𝒞; otherwise, a direct edge is added. This iterative process continues until all components are merged into a single weakly connected graph.

By prioritizing high-similarity connections and reusing existing ancestors where possible, this strategy ensures global connectivity with minimal additional nodes, preserving mutational interpretability and parsimony.

We model the MLG as a directed graph, where edges represent mutational transitions between variants. Directionality reflects the natural accumulation of mutations, but we also account for back mutations. To infer edge directionality in the MLG, we assign an edge *u* → *v* if the mutation set of variant *u* is a subset of that of *v*, i.e., *M* [*u*] ⊂ *M* [*v*], indicating a plausible forward mutation path. However, when the collection date of *v* precedes that of *u*, this contradiction may reflect either delays in collecting and sequencing or genuine evolutionary reversions, such as back mutations. Rather than disregarding such cases, we retain the directions based on collection dates and annotate the edges as potential reversion events.

#### Time Complexity

The overall time complexity of constructing the Mutation Learning Graph is driven by pairwise comparisons between mutation sets. Mutation profiles for all *k* variants are computed in 𝒪(*nk*) time, followed by pairwise edit distance and mutation similarity computations, each requiring 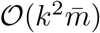, where 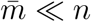 is the average number of mutations per sequence. Neighbor selection, tie resolution, and ancestor insertion contribute an additional 𝒪 (*k*^2^) in total. The final phase of component bridging, ensuring global graph connectivity, requires 𝒪 (*k* log *k*) time using max-heap and union-find operations. This scalable construction strategy allows us to handle large viral sequence datasets efficiently while preserving biological coherence.

### 2.7 Reconstruction of Inferred Ancestral Genomes

In our MLG, we introduced *inferred variants*, intermediate nodes deduced from mutational patterns. Each inferred variant was defined by its set of mutations relative to the reference genome (NC_045512.2). Genome alignments were reconstructed using their corresponding mutation sets in conjunction with the reference sequence, and the final genome sequences were obtained by eliminating all the alignment gaps.

### 2.8 MLG Data Set Structure

We propose a standardized dataset format for representing Mutation Learning Graphs (MLGs), designed to facilitate reproducible analysis and support a wide range of downstream learning tasks. Each dataset instance encodes both node features and edge features derived from the MLG, where nodes represent unique SARS-CoV-2 variants, either observed or inferred, and directed edges denote biologically plausible mutational transitions. The node features capture diverse genomic and temporal information, including sequence embeddings, mutation vectors, collection dates, and lineage annotations. The edge features encode mutational similarity, edit distance, mutation count, and evolutionary directionality. This structure enables biologically informed graph learning, providing a robust foundation for modeling viral evolution and predicting mutations. The node features and edge features are listed in Table 2 and 3 respectively.

**Table 2:**
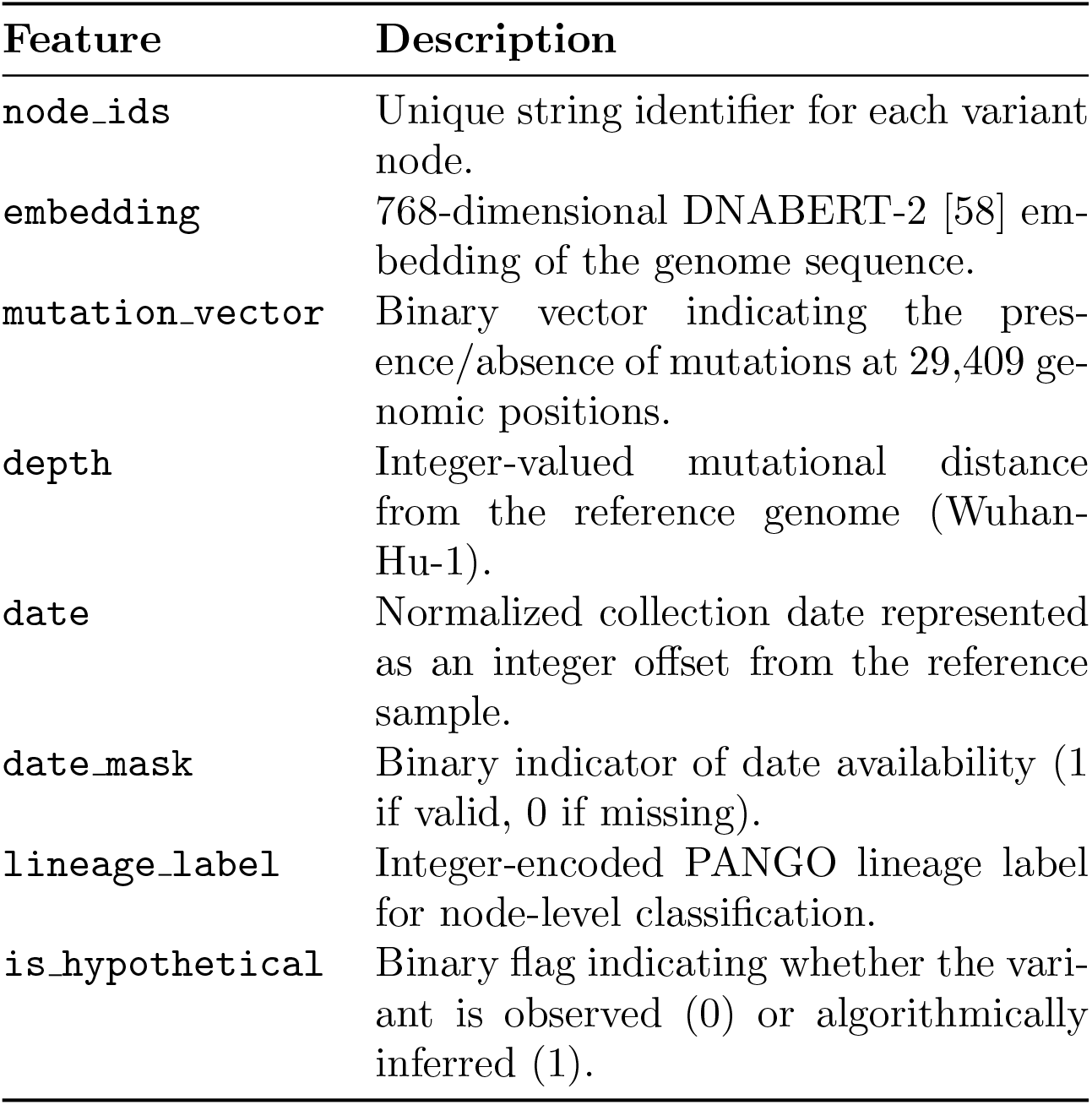
Node features in the MLG dataset.

| Feature | Description |
| --- | --- |
| `node_ids` | Unique string identifier for each variant node. |
| `embedding` | 768-dimensional DNABERT-2 [58] embedding of the genome sequence. |
| `mutation_vector` | Binary vector indicating the presence/absence of mutations at 29,409 genomic positions. |
| `depth` | Integer-valued mutational distance from the reference genome (Wuhan-Hu-1). |
| `date` | Normalized collection date represented as an integer offset from the reference sample. |
| `date_mask` | Binary indicator of date availability (1 if valid, 0 if missing). |
| `lineage_label` | Integer-encoded PANGO lineage label for node-level classification. |
| `is_hypothetical` | Binary flag indicating whether the variant is observed (0) or algorithmically inferred (1). |

**Table 3:** Edge features in the MLG dataset.

| Feature | Description |
| --- | --- |
| `source` | Integer index of the parent node. |
| `target` | Integer index of the child node. |
| `mutation_count` | Total number of mutations required to evolve from source to target. |
| `edit_distance` | Mutation-aware edit distance based on the differences in mutation sets. |
| `mutation_similarity` | Number of shared mutations between source and target variants. |
| `reverse` | Binary flag indicating reverse mutation (1 if reverse, 0 if forward). |

### 2.9 Problem Formulation and Benchmarking Tasks

To demonstrate the utility of our curated datasets for downstream graph-based learning, we benchmark two foundational tasks in the context of viral evolution modeling: node classification and link prediction. The node classification task serves as a *diagnostic benchmark*, verifying that the MLG representation captures lineage-related genomic and temporal patterns, and is not intended as a primary biological goal (Pangolin remains the current best practice for lineage assignment). In contrast, link prediction is biologically meaningful and directly aligned with one of our primary future research directions, forecasting plausible mutational transitions and unsampled future variants. While mutation frequency prediction may also be part of our future objectives, our focus in this work is to generate the MLG datasets as the foundational resource for such downstream forecasting tasks. These tasks validate that the curated MLG datasets are machine-learning-ready, directly compatible with graph neural networks without additional preprocessing, and capable of supporting both diagnostic analyses and biologically impactful prediction objectives.

#### Algorithm 1 Ancestor-Joining^1^

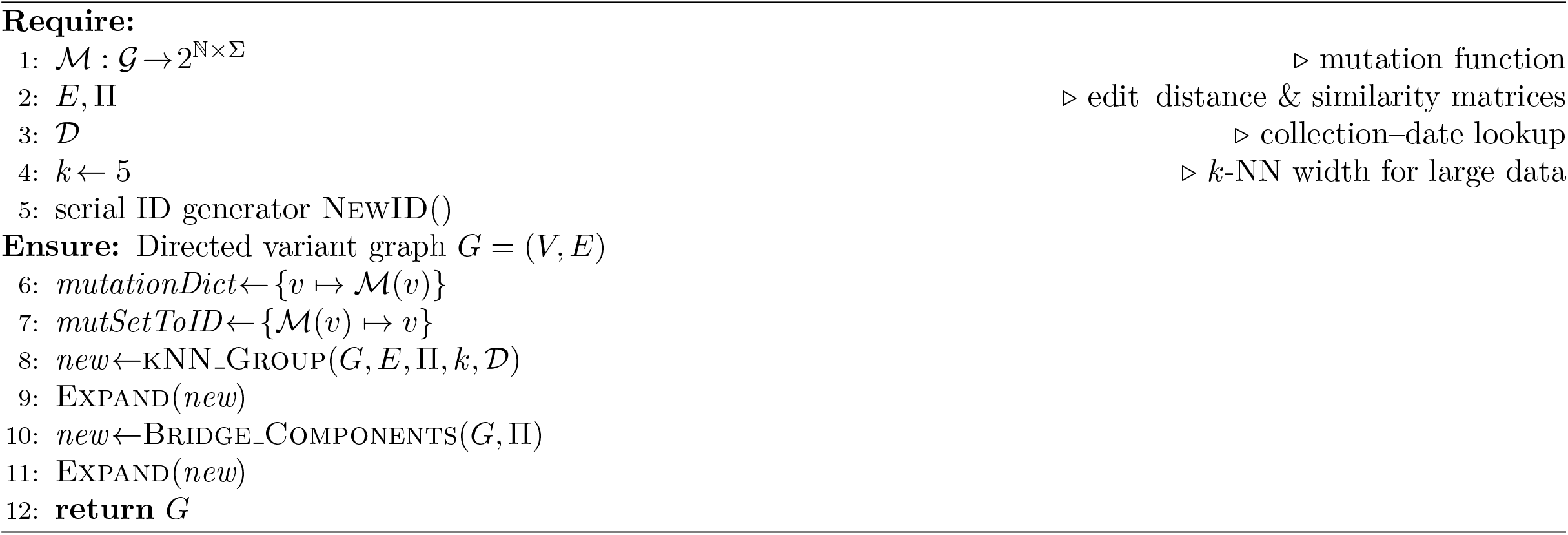

#### 2.9.1 Node Classification: Inferring Lineage

We formulate lineage classification as a node-level supervised learning task on an MLG, *G* = (*V, E*), where nodes *v* ∈ *V* represent SARS-CoV-2 variants (observed and inferred) and edges (*u, v*) ∈ *E* indicate plausible mutational transitions. Each node is associated with a feature vector *x*_*i*_ ∈ ℝ^*d*^, as described in section 2.8, encompassing sequence embeddings, mutation vectors, and relevant metadata.

The objective is to learn a function *f* : ℝ^*d*^→𝒴, where 𝒴 denotes the set of Pango [38] lineage labels, under a multi-class classification framework. Only nodes corresponding to *observed* variants (i.e., `is_hypothetical` = 0) are used for training, validation, and testing. *Inferred* intermediate nodes are excluded from supervision, but they participate in message passing during training. This problem formulation enables benchmarking of GNN-based models on their ability to classify viral lineages using both genomic and topological signals embedded in the MLG.

#### 2.9.2 Baseline Models for Node Classification

We evaluated five baseline models for node classification: GCN [26], GraphSAGE [17], GAT [48], GGNN [31], and a Multi-Layer Perceptron (MLP) [40] as a feature-only baseline. All models were implemented using PyTorch Geometric and trained over 200 epochs with the Adam optimizer, using a learning rate of 0.01 (except GAT with 0.005) and weight decay of 5 *×* 10^*−*4^. The input feature matrix included DNABERT-2 embeddings, PCA-transformed mutation vectors, sequencing depth, normalized collection date, and a binary indicator for hypothetical variants. Node classification was restricted to observed variants with known lineage labels, stratified into 3-fold train/validation/test splits. Inferred nodes were retained in the graph for message passing but excluded from supervision. To mitigate class imbalance, we employed class-weighted cross-entropy loss, based on the label distribution of each fold. Evaluation was conducted using 10 bootstrap trials, reporting mean ± standard deviation for accuracy, F1 score, AUROC, and AUPRC.

#### 2.9.3 Link Prediction: Inferring Mutational Transitions

In the edge prediction task, the goal is to infer plausible mutational transitions between variant pairs. Let *G* = (*V, E*) denote the directed MLG, where each edge (*u, v*) ∈ *E* corresponds to a mutational transition from variant *u* to variant *v*. The task is to learn a scoring function *f* (*u, v*) ∈ [0, 1] that predicts whether an edge should exist between two nodes. This problem is framed as a binary classification problem: existing edges are positive samples, and an equal number of non-edges (excluding self-loops) are sampled as negatives.

#### 2.9.4 Baseline Models for Link Prediction

We evaluated four representative graph neural network architectures for the link prediction task: GraphSAGE [17], GAT [48], GGNN [31], and VGAE [27], along with a non-graph baseline using a multilayer perceptron (MLP) [40]. All models integrate node-level features (e.g., DNABERT-2 embeddings, PCA-transformed mutation vectors, depth, temporal metadata) and edge-level attributes (e.g., mutation count, edit distance, mutation similarity, directionality).

Graph-based models employ a graph encoder to produce node embeddings, followed by an MLP decoder for edge classification. In GraphSAGE and GAT, node embeddings are concatenated with edge features, with GAT using multi-head attention for neighborhood aggregation. GGNN applies gated recurrent units for iterative message passing, while VGAE uses a probabilistic GraphSAGE encoder and variational MLP decoder with binary cross-entropy and KL-divergence loss. The MLP baseline omits the graph encoder, directly concatenating raw node features of each endpoint with edge attributes for classification, providing a reference for the added value of structural learning. All models were trained using stratified splits and 10-fold bootstrapping to address class imbalance.

## 3 Results

### 3.1 Lineage Distribution Across Data Sets

We analyzed the lineage distribution in each of the regional SARS-CoV-2 data sets to assess class balance and diversity for downstream classification tasks. A summary of key statistics, including the count of distinct lineages, the dominant lineage, its proportion, and Shannon entropy, is provided in Table 4. This table reveals substantial variation across data sets. For example, Nigeria and Bangladesh are dominated by a single lineage (AY.36 and B.1.617.2, respectively), whereas China and Queensland exhibit a more balanced lineage spread with higher entropy scores.

**Table 4:**
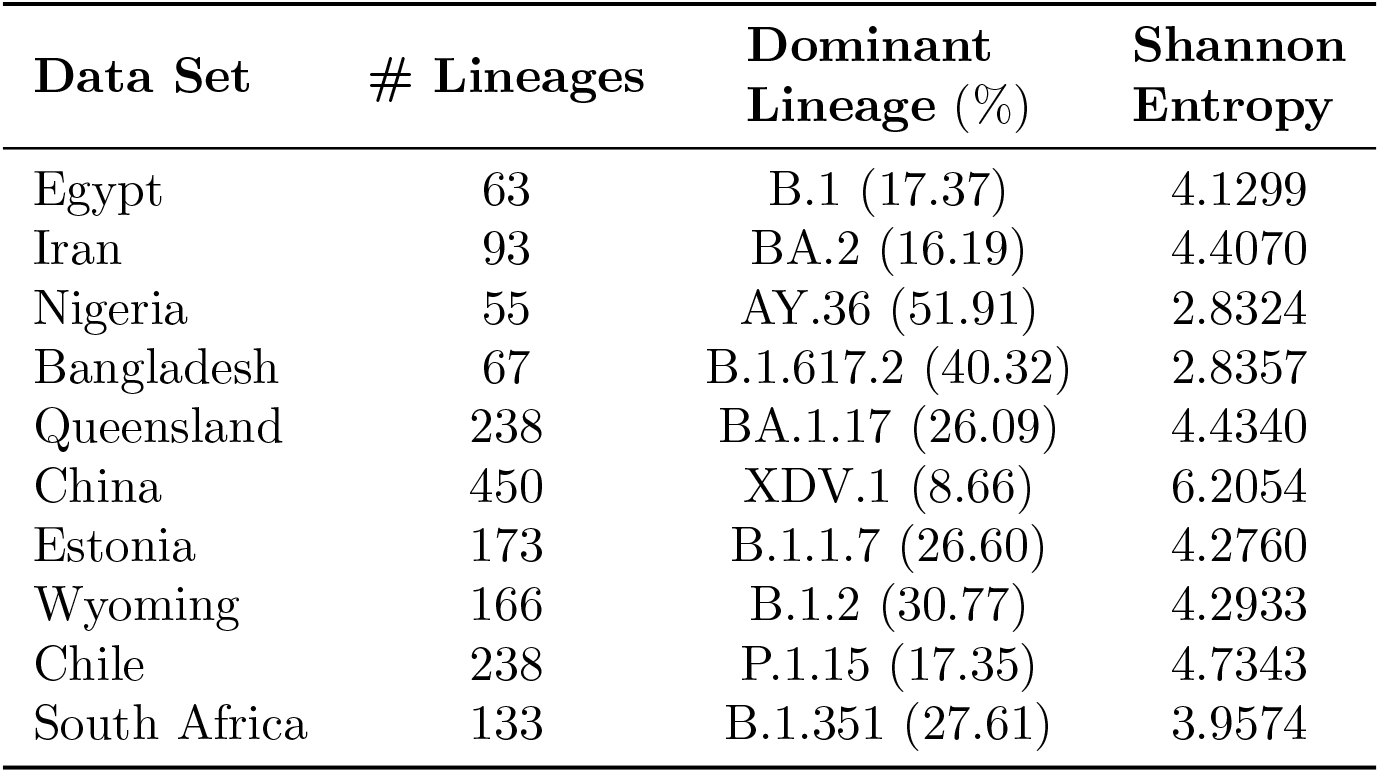
Summary statistics of lineage distributions across SARS-CoV-2 data sets.

To visualize lineage diversity, Fig. 3 presents violin plots of lineage distributions across all ten datasets. The shape of each violin reflects the density of lineage frequencies, sharp peaks (e.g., Nigeria, Bangladesh) indicate dominance by a few lineages. In contrast, broader shapes (e.g., China, Estonia) suggest greater diversity. These patterns highlight regional differences in viral evolution and sampling, underscoring the importance of evaluating models under diverse epidemiological conditions. Detailed lineage breakdowns are provided as bar charts in the Supplementary Material, enabling a fine-grained understanding of lineage composition in each dataset.

**Figure 3:**
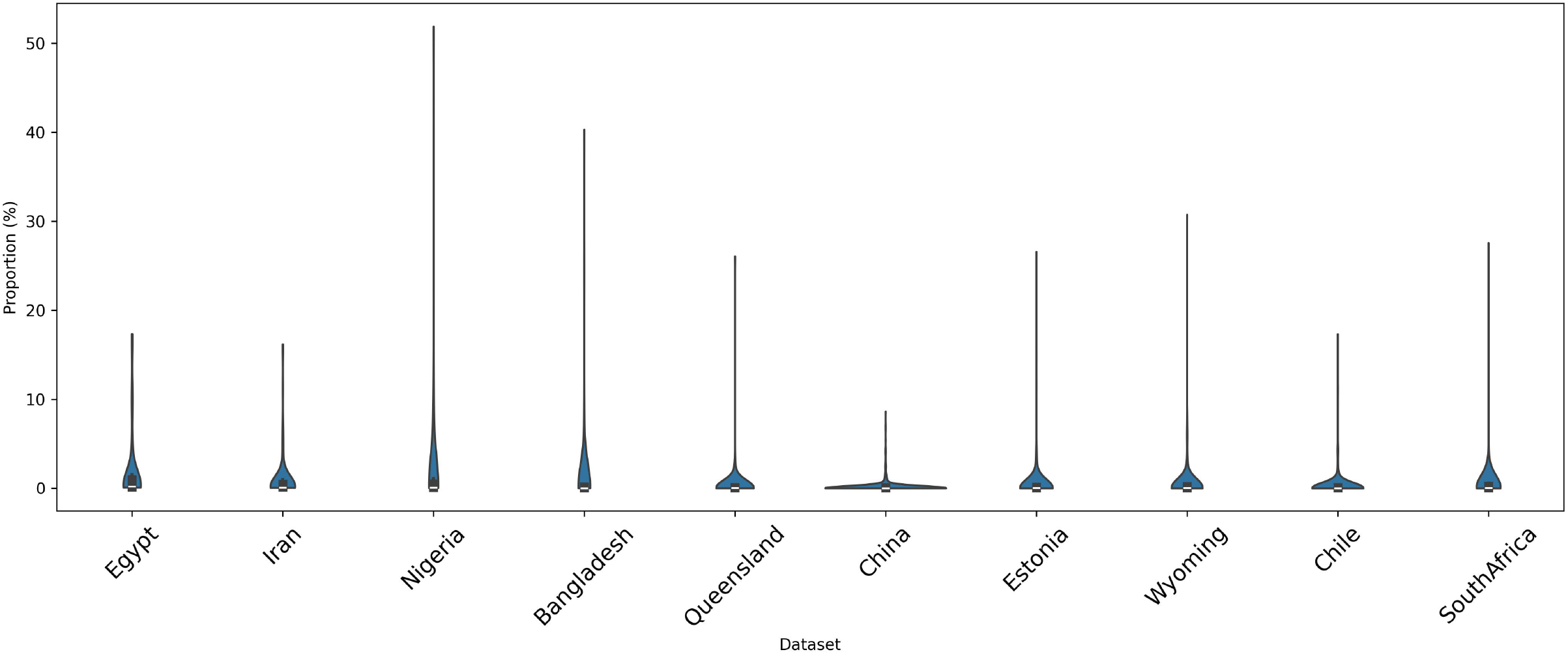
Violin plot showing the distribution of lineage proportions across ten SARS-CoV-2 regional data sets.

### 3.2 Validation of the Inferred Variants

To assess the biological plausibility of these inferred sequences, we utilized Pangolin [38] v4.3.0, along with the latest pangoLEARN model, for lineage assignment. All inferred variant genomes were successfully validated, with no ambiguity reported in any data set (i.e., all `scorpio_call` and `ambiguity_score` fields were `NaN`, all `qc_status` were `pass`). In some cases, lineage was assigned directly from a designation hash, which indicates an exact mutation-level match between the reconstructed variant and an existing SARS-CoV-2 lineage, i.e., a representative sequence for that lineage. These inferred nodes, functioning as internal nodes in the MLG, represent evolutionarily credible intermediates that connect observed genotypes along valid mutational trajectories.

Confident phylogenetic placements from UShER (e.g., B(1/1)) confirmed that all inferred nodes aligned well with the global SARS-CoV-2 tree. Minor conflicts (e.g., conflict scores of 0.5) were observed in a few cases, consistent with their role as inferred intermediates rather than directly sampled variants. These partial mismatches suggest plausible evolutionary connections rather than misclassification. Table 5 summarizes the lineage assignments, showing that all inferred nodes were assigned valid SARS-CoV-2 lineages, primarily within lineage B. Full assignment details, including UShER placements and conflict scores, are available in the Supplementary Material.

**Table 5:**
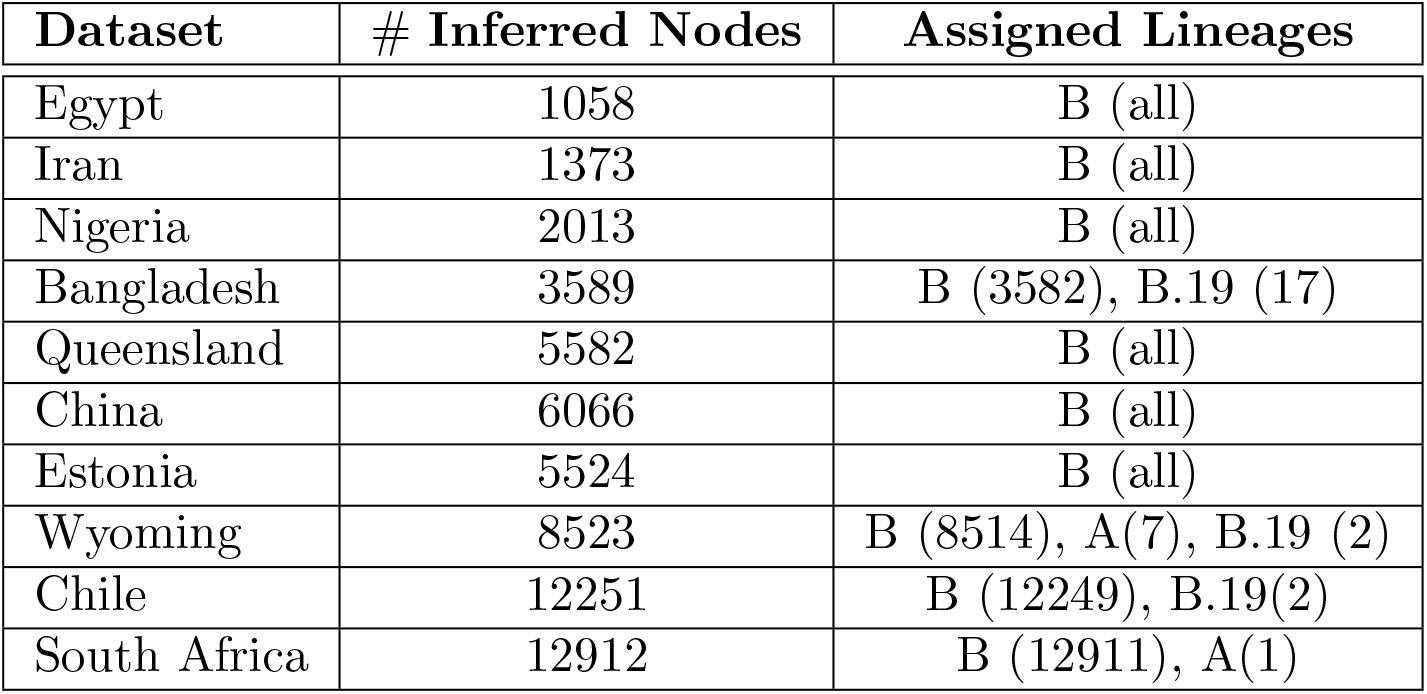
Summary of Pangolin-based lineage assignment to the inferred nodes across datasets.

| Dataset | # Inferred Nodes | Assigned Lineages |
| --- | --- | --- |
| Egypt | 1058 | B (all) |
| Iran | 1373 | B (all) |
| Nigeria | 2013 | B (all) |
| Bangladesh | 3589 | B (3582), B.19 (17) |
| Queensland | 5582 | B (all) |
| China | 6066 | B (all) |
| Estonia | 5524 | B (all) |
| Wyoming | 8523 | B (8514), A(7), B.19 (2) |
| Chile | 12251 | B (12249), B.19(2) |
| South Africa | 12912 | B (12911), A(1) |

### 3.3 SNP Distribution Analysis

To explore the mutational landscape at the single-nucleotide level, we analyzed the distribution of all twelve possible single-nucleotide polymorphisms (SNPs) across our ten region-specific SARS-CoV-2 datasets. Fig. 4 presents a grouped bar chart showing the percentage of each SNP type per data set.

**Figure 4:**
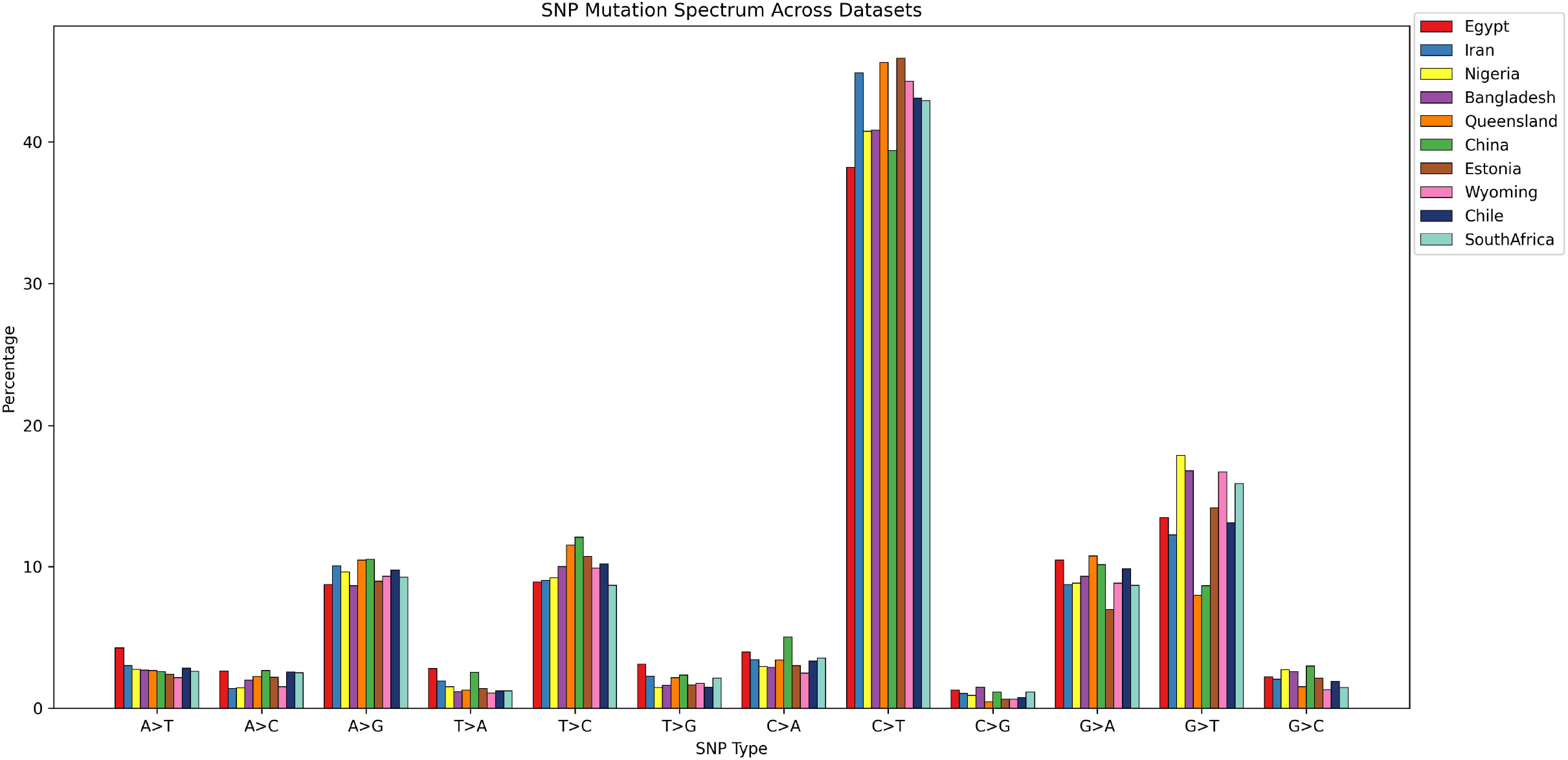
Barchart of the SNP distribution in the data sets.

We observe a clear predominance of **C** → **T** transitions across all datasets, followed by **G** → **T** and **G** → **A** substitutions. This trend is consistent with previously reported global patterns of SARS-CoV-2 mutation [35, 55, 15], where **C**→**T** transitions are the most frequent mutation class, likely driven by host-mediated RNA editing mechanisms such as APOBEC and ADAR activity [16]. The consistency of these patterns across data sets supports the biological plausibility of our mutation modeling approach.

### 3.4 Genomic Positions of Mutations

Understanding the genomic positions of mutations is essential for characterizing how viral genomes evolve. Mutations do not occur uniformly across the genome, and different geographical regions exhibit distinct patterns in the locations and frequencies of these mutations, as shown in Fig. 5. Founder effects, selective pressures, and transmission dynamics often shape these region-specific trends. Epidemiologically, tracing such mutation patterns is vital for identifying evolutionary hotspots. To reflect these spatial and temporal trends, our datasets are constructed separately for each geographic location, preserving the distinct mutational signatures specific to that region.

**Figure 5:**
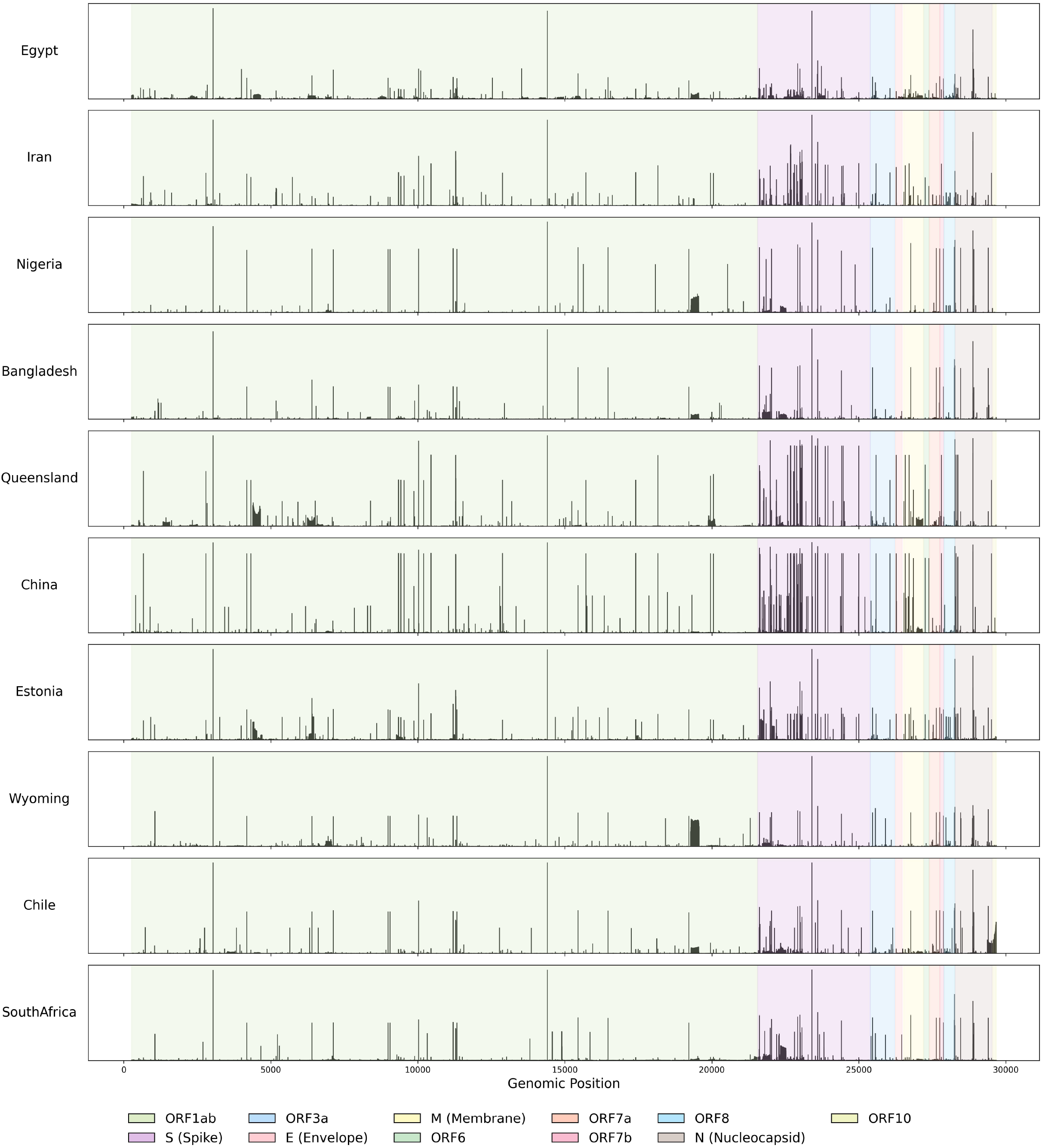
Percentages of mutation frequencies across SARS-CoV-2 genomic positions for all the data sets.

### 3.5 Summary Statistics of MLGs

We constructed Mutation Learning Graphs (MLGs) for ten geographically diverse SARS-CoV-2 datasets. Table 6 summarizes the structural properties of the resulting graphs, including the number of observed and inferred nodes, total edges, average node degree, graph density, maximum depth from the reference node, and the proportion of nodes with at least one outgoing edge.

**Table 6:** Summary statistics of Mutation Learning Graphs (MLGs) across 10 datasets.

| Dataset | # Observed | # Inferred | # Edges | Avg Degree | Graph Density | Max Depth | % Out-Degree $\geq 1$ | Reverse Edges |
| --- | --- | --- | --- | --- | --- | --- | --- | --- |
| Egypt | 898 | 1058 | 2775 | 2.84 | 0.00073 | 6072 | 61.96 | 2.56 |
| Iran | 1285 | 1373 | 4166 | 3.13 | 0.00059 | 507 | 56.81 | 0.62 |
| Nigeria | 1934 | 2013 | 6090 | 3.09 | 0.00039 | 346 | 57.97 | 1.12 |
| Bangladesh | 3125 | 3589 | 10309 | 3.07 | 0.00023 | 1828 | 58.52 | 0.88 |
| Queensland | 5427 | 5582 | 16627 | 3.02 | 0.00014 | 2863 | 61.10 | 1.58 |
| China | 7140 | 6066 | 20283 | 3.07 | 0.00012 | 1102 | 56.53 | 2.01 |
| Estonia | 10494 | 9946 | 31302 | 3.06 | 0.00008 | 1651 | 57.43 | 1.73 |
| Wyoming | 11164 | 10929 | 34030 | 3.08 | 0.00007 | 1803 | 57.41 | 1.50 |
| Chile | 14131 | 11826 | 39329 | 3.03 | 0.00006 | 1551 | 57.76 | 1.50 |
| South Africa | 12055 | 10288 | 34351 | 3.08 | 0.00007 | 1917 | 58.34 | 1.73 |

Across all datasets, the MLGs exhibit relatively low graph density despite incorporating thousands of unique sequences. This sparsity reflects the edge construction strategy, which employs a *k*-nearest neighbor approach to connect nodes based on mutational similarity and temporal precedence, rather than considering all nearest neighbors. This heuristic was selected to ensure computational scalability for large datasets, where evaluating all pairwise mutational relationships would be infeasible.

### 3.6 Cross-Regional MLG Comparison

To quantify structural differences between MLGs from different regions, we computed the Jensen–Shannon divergence (JSD) between normalized edge feature distributions. Figure 6 shows the JSD matrix for *mutation similarity*, which exhibits the strongest divergence patterns, particularly for pairs such as Egypt–Estonia and Egypt–Nigeria. This indicates substantial variation in mutational connectivity profiles across regions. JSD analyses for *edit distance* and *mutation count* show more moderate differences and are included in the Supplementary Material.

**Figure 6:**
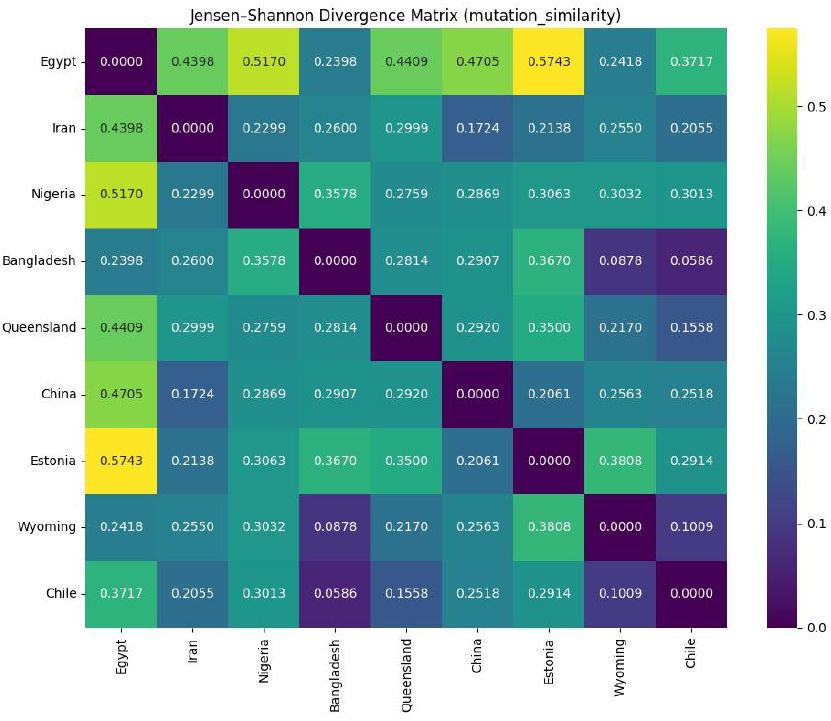
Jensen–Shannon divergence matrix for *mutation similarity* across ten regional MLGs. Higher values indicate greater dissimilarity in mutational connectivity patterns.

We further examined the in-degree and out-degree distributions of all regional MLGs to characterize differences in network topology. Degree counts were normalized and plotted on log-log scales in a combined panel (Fig. 7). Both distributions exhibit heavy-tailed patterns, consistent with preferential attachment processes in viral evolution. While the overall shapes are similar across regions, variations in slope and tail weight reflect differences in mutational sourcing and propagation between datasets. Individual plots for in-degree and out-degree distributions are provided in the Supplementary Material.

**Figure 7:**
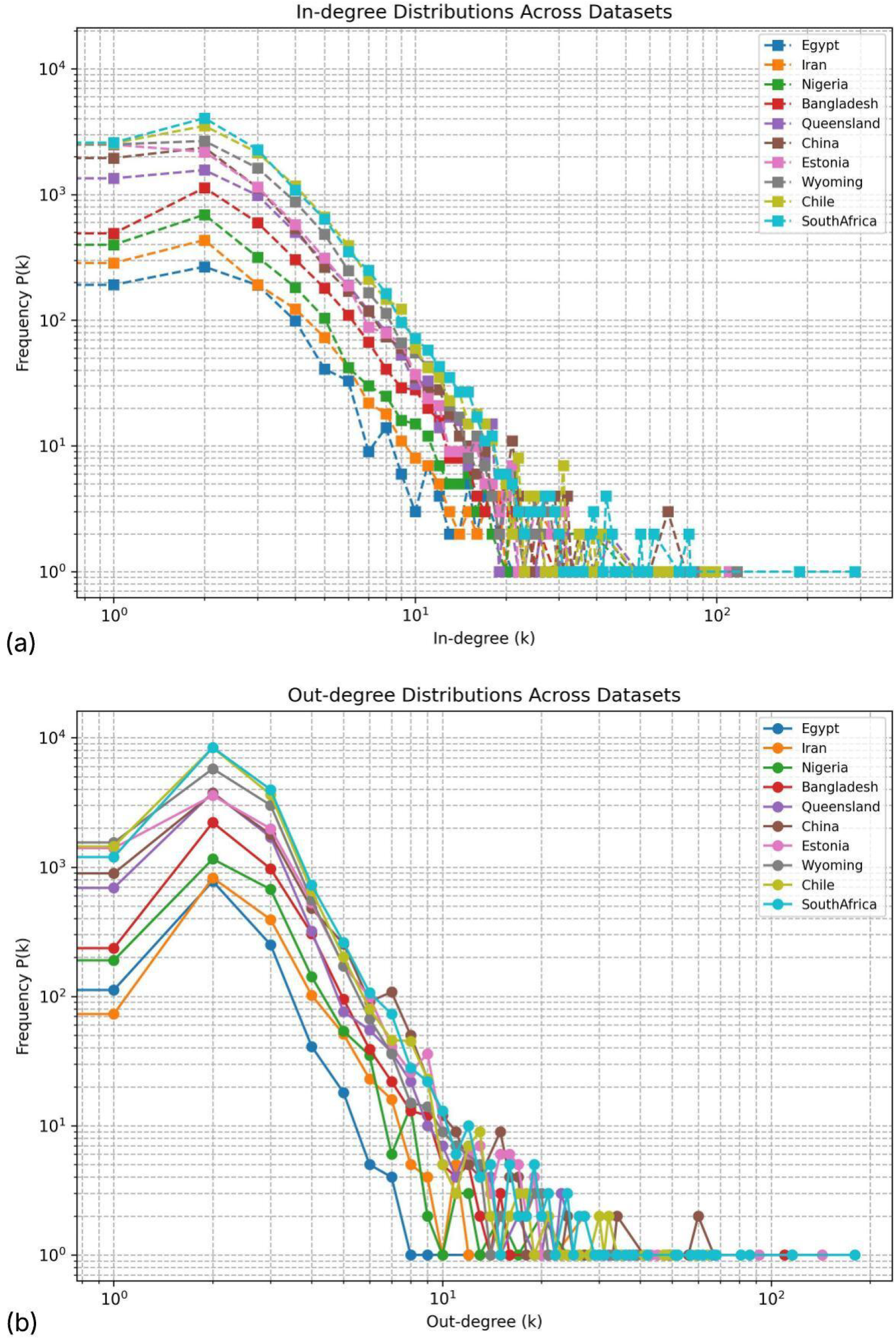
In-degree and out-degree distributions across datasets. Each curve corresponds to a regional MLG, with degree counts plotted on log-log scales.

### 3.7 Reverse Mutation

In addition to spatial information, we include the collection dates of observed variants to capture temporal patterns in viral evolution. These dates are used to infer edge directionality along with the mutation sets of the variant nodes, and they play a crucial role in identifying potential back mutations. By incorporating both the genomic context and sampling timeline, our data sets support a more biologically grounded modeling of MLGs.

While most mutational transitions in the network proceed in a forward direction, accumulating changes from ancestral to descendant variants, we also observe a subset of edges that appear to reverse this trajectory. Specifically, we define an edge as a *reverse mutation* when variant *u* has a mutation set ℳ [*u*] ⊂ ℳ [*v*] but was collected after *v*, thus reversing the expected temporal order (described in section 2.6). Although such events may result from either reverse mutations or delays in sample collection or sequencing, we treat them as biologically potential reverse mutations. This strategy enables the models to learn from both canonical forward mutation events and rare but biologically significant reversions, enhancing the fidelity of the resulting MLG.

To quantify their prevalence, we calculated the proportion of reverse mutations in each dataset (Table 6). While forward mutation remains dominant across all datasets, a consistent proportion (1% to 3% of the total) of potential reverse mutation edges is observed.

### 3.8 Benchmarking Graph-Based Learning Tasks

#### 3.8.1 Node Classification Performance

As shown in Fig. 8, node classification performance varies substantially across datasets and models, highlighting the influence of network size, lineage imbalance, and feature quality. GraphSAGE and GGNN consistently achieved strong accuracy and F1 scores across most datasets, particularly in Nigeria, Bangladesh, and Estonia. These datasets feature larger variant pools and moderately diverse lineage distributions, enabling more stable graph-based learning. The MLP baseline, which does not utilize graph structure, performed well in accuracy and F1 for several datasets (e.g., Egypt and Iran), indicating that DNABERT-2 embeddings and mutation vectors carry strong lineage signals. Its low *macro*-AUPRC is largely due to the metric’s sensitivity to class imbalance, as absent rare lineages in the test split are assigned AP = 0, lowering the macro average even when predictions for present classes are accurate. GAT exhibited inconsistent results, likely due to its reliance on attention mechanisms that are sensitive to sparsity and class imbalance. Notably, AUPRC values were more discriminative than accuracy or F1, revealing the impact of long-tailed lineage distributions. AUROC was omitted from the heatmap because it could only be stably computed for GCN; other models produced NaN values due to prediction collapse or extreme class imbalance. Numerical details are included in the Supplementary Material.

**Figure 8:**
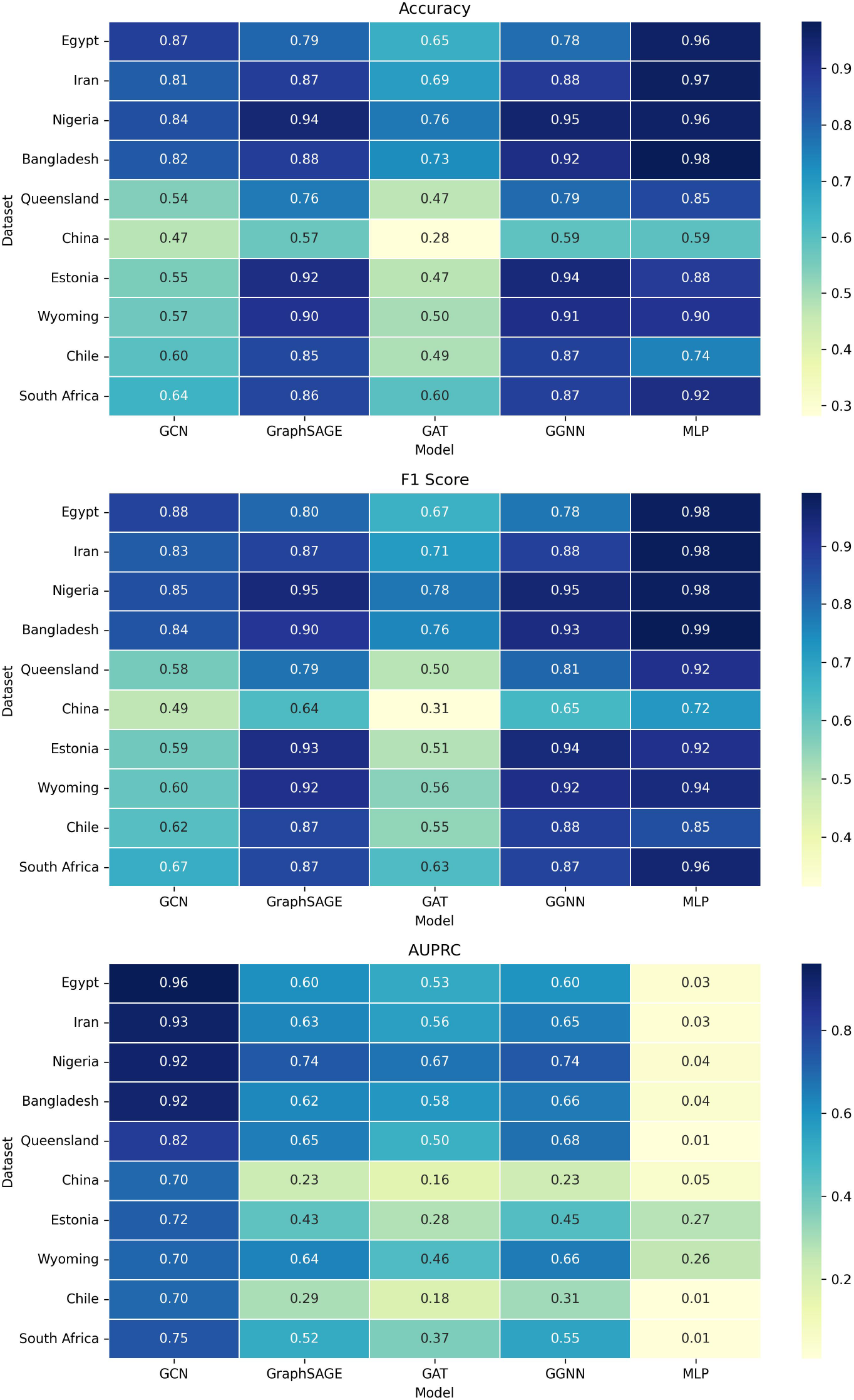
Heatmaps showing node classification performance across ten SARS-CoV-2 datasets using five baseline models (GCN, GraphSAGE, GAT, GGNN, and MLP).

#### 3.8.2 Link Prediction Performance

#### 3.8.3 Link Prediction Performance

Fig. 9 presents heatmaps of model performance across four metrics — Accuracy, F1 Score, AUROC, and AUPRC — for the edge prediction task on the ten datasets. GraphSAGE and VGAE consistently outperformed other methods across datasets. GraphSAGE achieved the highest average AUROC, while VGAE led in AUPRC, reflecting its strength in modeling imbalanced mutational transitions. The success of VGAE is attributable to its variational framework and generative edge reconstruction, which proved robust across datasets of varying sizes and lineage complexities (e.g., China, Bangladesh, and Estonia). GGNN achieved the highest accuracy in several datasets, including those from Iran, Bangladesh, and South Africa, but exhibited substantial drops in F1 Score, indicating sensitivity to edge class imbalance and possible overfitting to dominant mutation pathways. GAT performed poorly across all metrics, suggesting that its attention mechanisms are less effective under the sparsity and heterogeneity of viral mutation networks.

**Figure 9:**
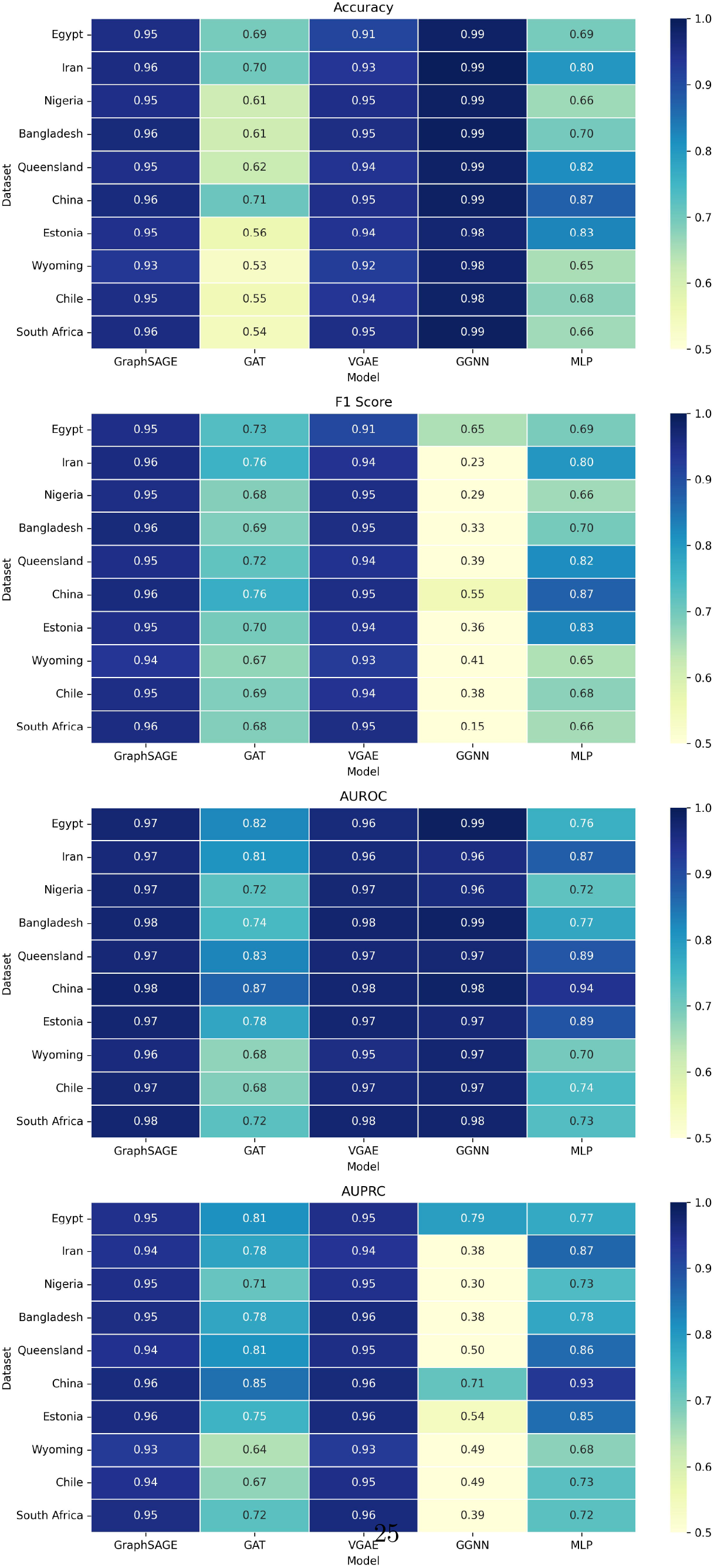
Heatmaps showing edge prediction performance across ten SARS-CoV-2 datasets using four baseline models (GraphSAGE, GAT, VGAE, GGNN, and MLP).

The non-graph MLP baseline, which classifies edges directly from concatenated node and edge features without leveraging graph structure, consistently underperformed the GNN-based models. While MLP attained moderate scores in datasets with strong attribute signals (e.g., China, Queensland, and Iran), it lagged significantly on others, underscoring the added predictive value of structural context learned by graph encoders.

A notable challenge in this task arises from a small proportion (1%–3%) of biologically plausible but topologically inconsistent edges that represent reverse mutations. These edges, caused by either true reverse mutation or sampling delays, conflict with the forward-only mutation pattern and can introduce label noise. Despite this, both GraphSAGE and VGAE maintained strong predictive performance, demonstrating their suitability for capturing signals of mutational transitions. These results highlight the utility of graph-based models for evolutionary inference and validate the proposed MLG framework for realistic modeling of mutation transitions. A comprehensive summary of all numerical results is provided in the Supplementary Material.

## 4 Discussion

In this work, we introduced the Mutation Learning Graph, a curated graph-based dataset framework that captures fine-grained mutational relationships among SARS-CoV-2 variants across diverse regions. By integrating observed and inferred variants and encoding positional mutation data, MLG provides a biologically grounded platform for studying viral evolution.

We evaluated MLGs through two benchmark tasks. Our use of node classification serves primarily as a diagnostic task to assess the richness of encoded features and lineage signals in the graph. While not a direct objective of this study, it confirms that the MLG representation supports meaningful learning across variant-level genomic and temporal features. Link prediction, on the other hand, is more biologically aligned with our goals, as it corresponds to modeling mutational transitions and evolutionary trajectories between variants. Through benchmarking, we identified models such as GraphSAGE and VGAE that consistently outperform others across diverse mutation landscapes, providing valuable guidance for choosing architectures suited to viral genomic data. Notably, VGAE not only excels in reconstructing known transitions but also offers a reusable generative layer for predicting plausible, unobserved mutational links, eliminating the need to recompute complete similarity or distance matrices and enabling scalable mutation forecasting.

A central component of MLG construction is the Ancestor-Joining algorithm, which introduces inferred variants to maintain consistent mutational propagation. These inferred nodes, validated by lineage assignments, contribute to coherent graph topology and possibly represent unsampled intermediates.

Despite its strengths, the current version of MLG remains a sparse representation due to the computational constraints of the k-nearest neighbor algorithm used during edge construction. As a result, all plausible mutational paths are not present, potentially limiting downstream tasks. To address this, a key future direction involves enriching the MLG by leveraging VGAE-based link prediction to predict plausible but unobserved evolutionary transitions and add missing edges to the graph. This generative extension would enhance the structural completeness of the graph and enable mutation forecasting, a crucial step toward predicting future variants.

Finally, observed lineage imbalance, regional mutation spectra, and reverse mutations emphasize the need for location-aware and mutation-centric modeling approaches. We release all MLG datasets and reproducible code to support further research into viral evolution and prediction.

## 5 Conclusion

We presented the Mutation Learning Graph (MLG), a scalable and interpretable framework for modeling viral evolution through graph-based learning. By organizing observed and inferred SARS-CoV-2 variants into a mutation-informed graph enriched with genomic and temporal features, MLG captures the mutational dynamics across ten geographically diverse regions. The resulting datasets reflect real-world lineage heterogeneity, positional mutation trends, and plausible intermediate variants, offering a rich foundation for downstream modeling.

Through benchmarking on node classification and link prediction tasks, we demonstrated that while MLG supports meaningful learning, general-purpose GNNs often struggle with challenges such as lineage imbalance, reverse mutations, and regional diversity. Our results underscore the need for mutation-aware, region-specific models that are tailored to the complexities of viral evolution.

While our MLG does not claim to represent the exact evolutionary history of SARS-CoV-2, it provides a mutation-informed structural organization of variant genomes. This abstraction captures key dimensions of viral diversity, including genome sequence changes, mutation profiles, mutational distances, and transition pathways, offering a multifaceted perspective on variant relationships. In doing so, it facilitates both predictive modeling and hypothesis generation for future evolutionary outcomes. Looking ahead, we aim to enrich MLG by leveraging generative models, particularly VGAE, for predicting plausible edges that are missed due to computational constraints, thereby enabling scalable mutation forecasting.

Our framework is broadly applicable to other viral outbreaks where sequence data are available. By releasing reproducible code and curated datasets, we enable the community to build and extend predictive models for viral evolution, variant emergence, and outbreak preparedness.

---

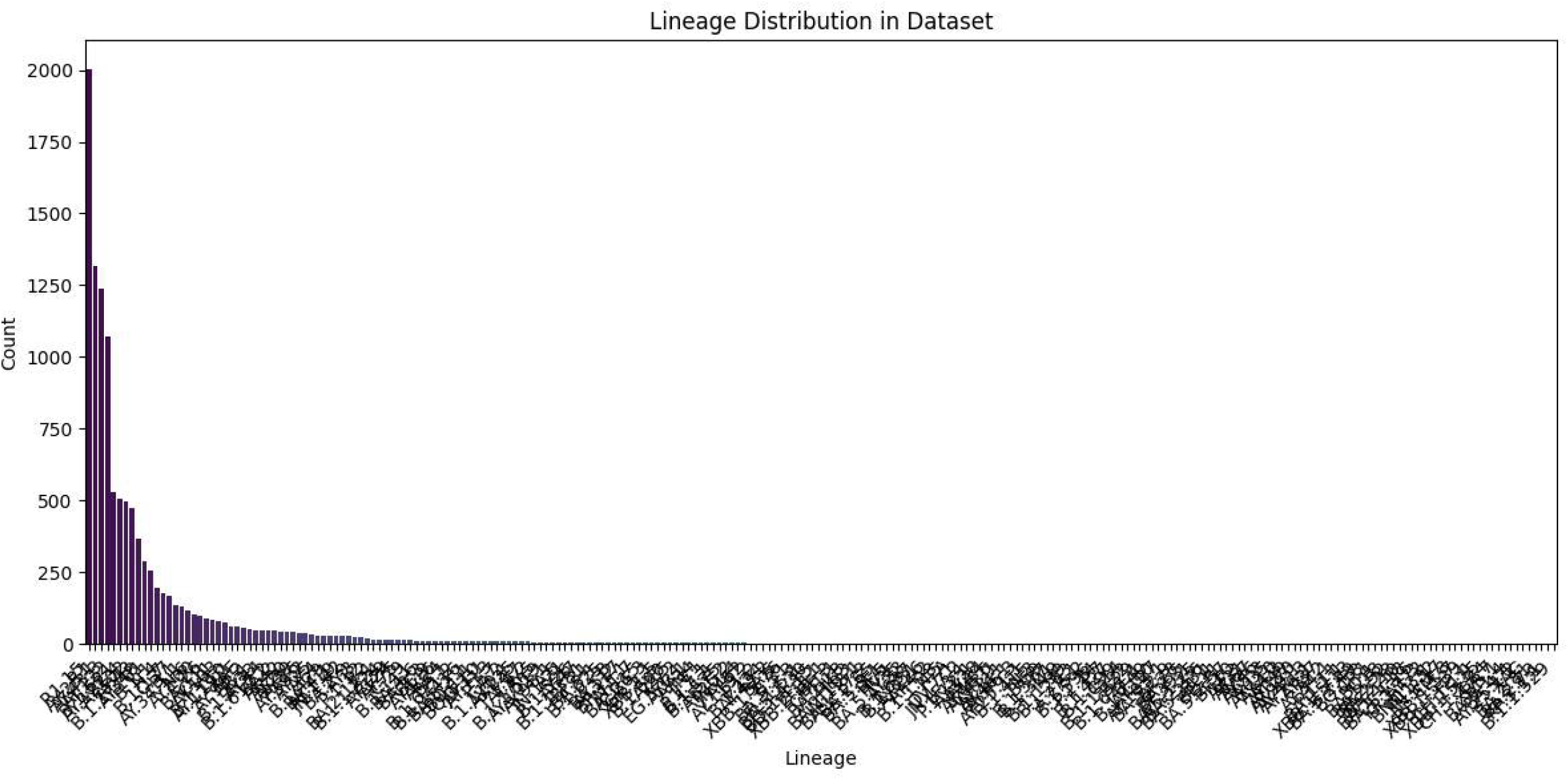

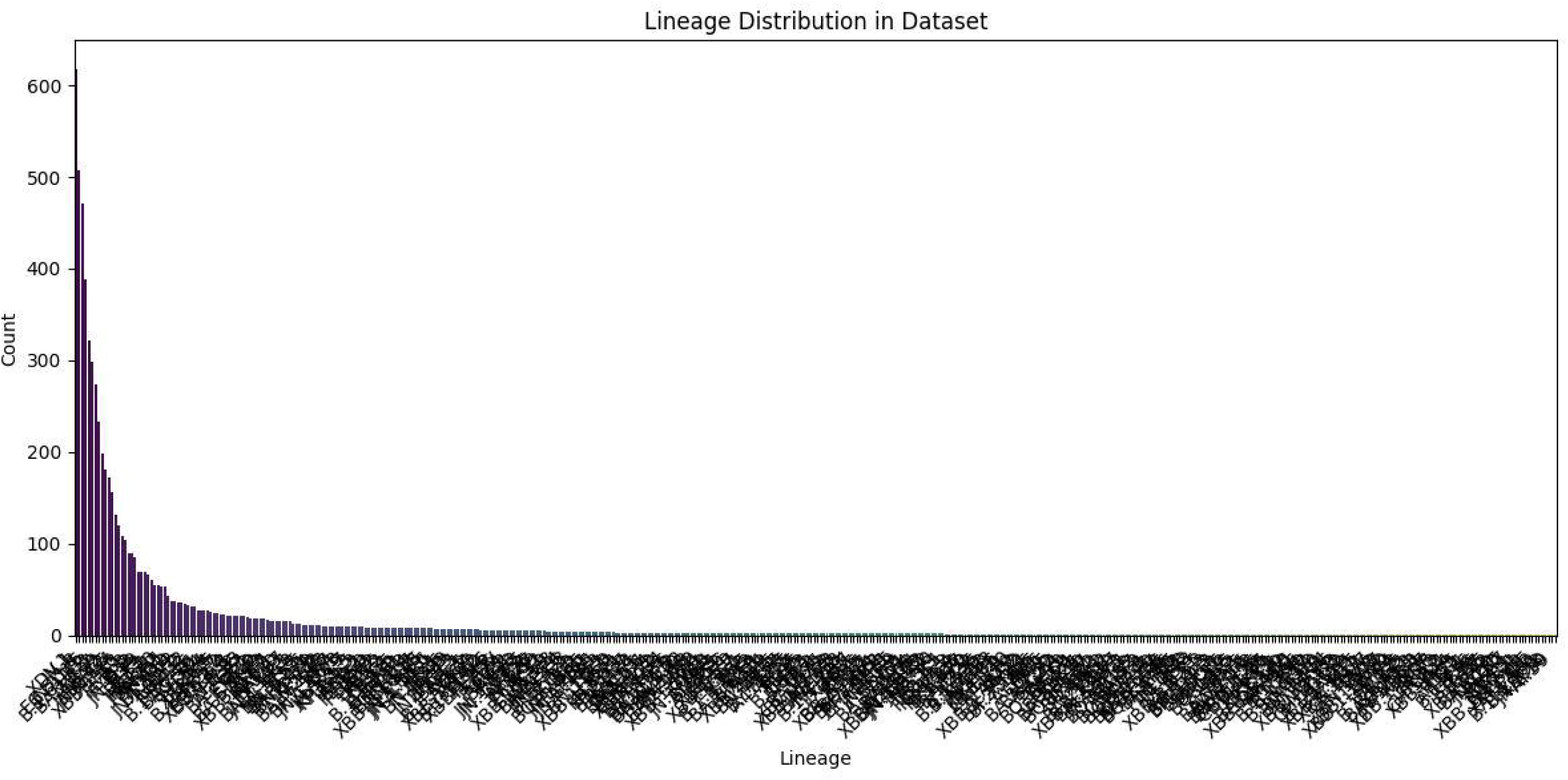

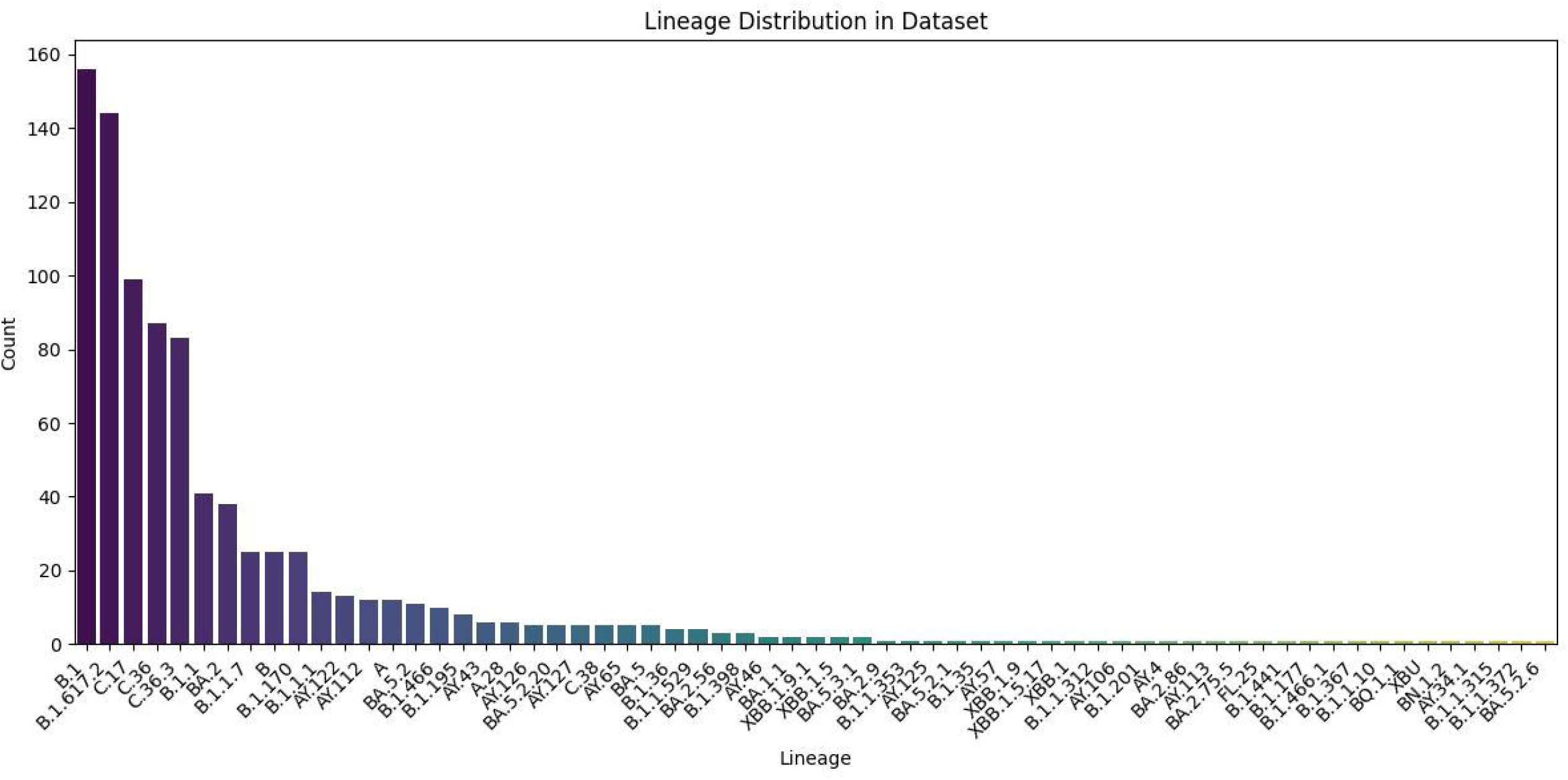

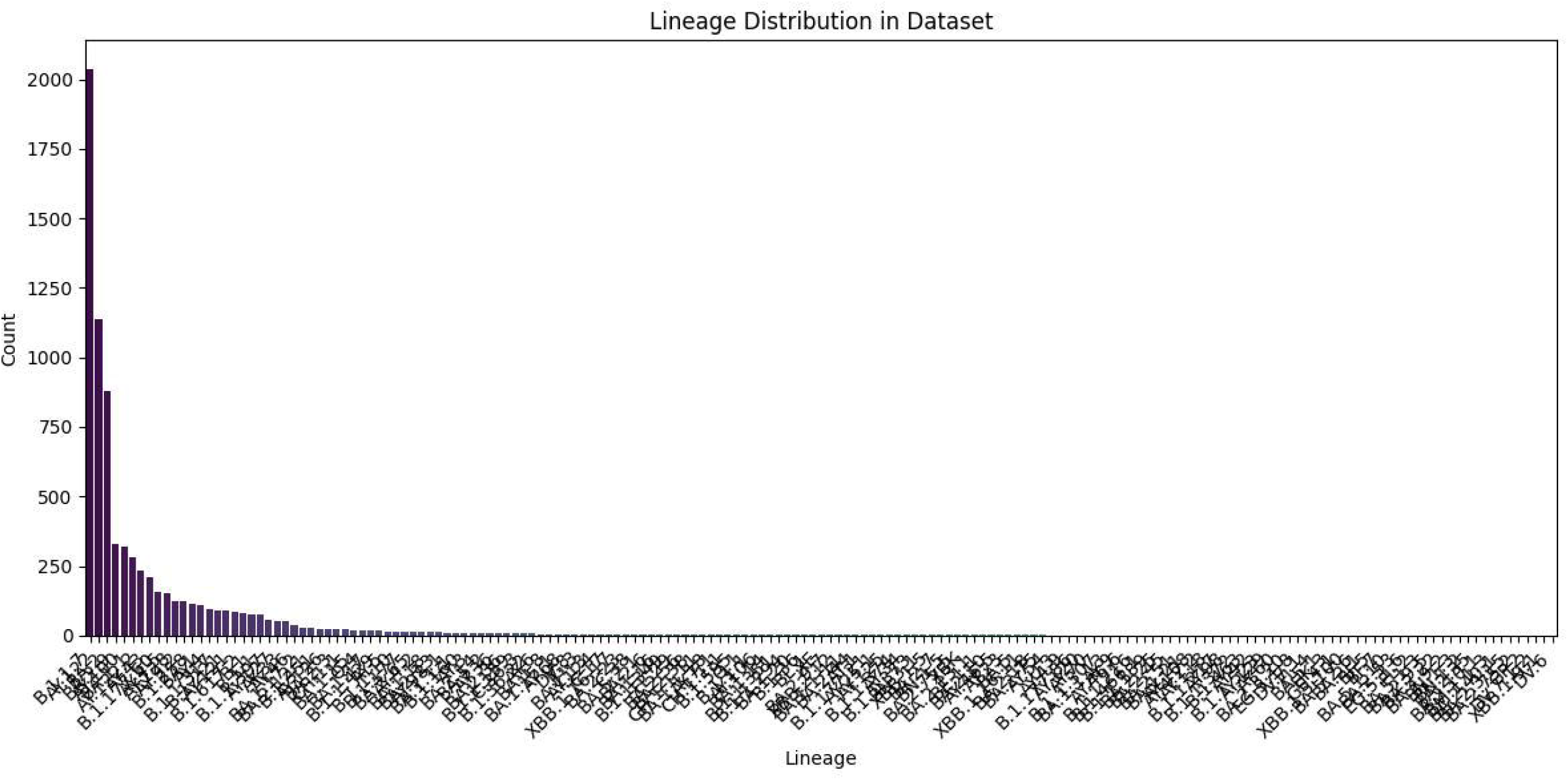

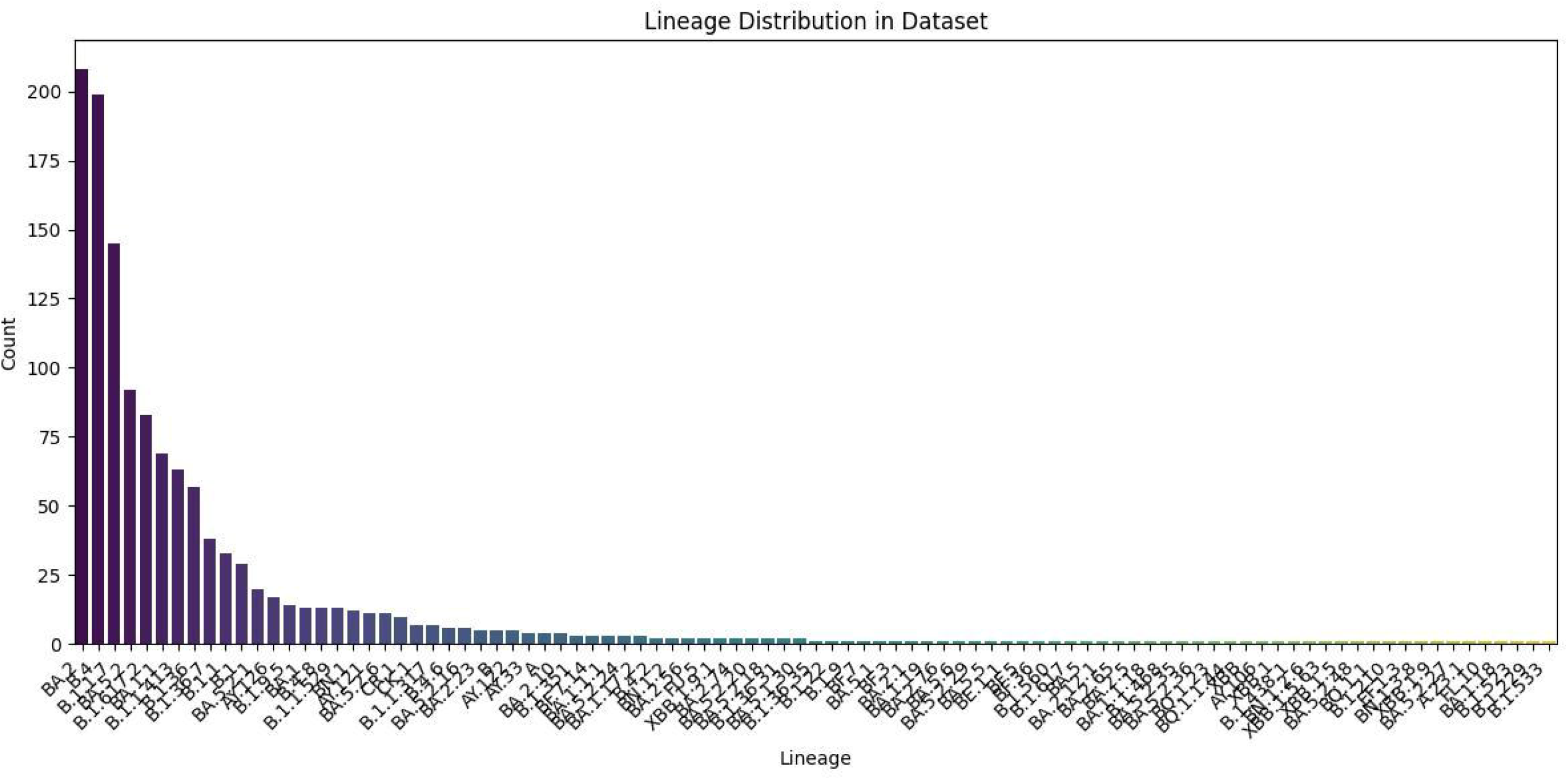

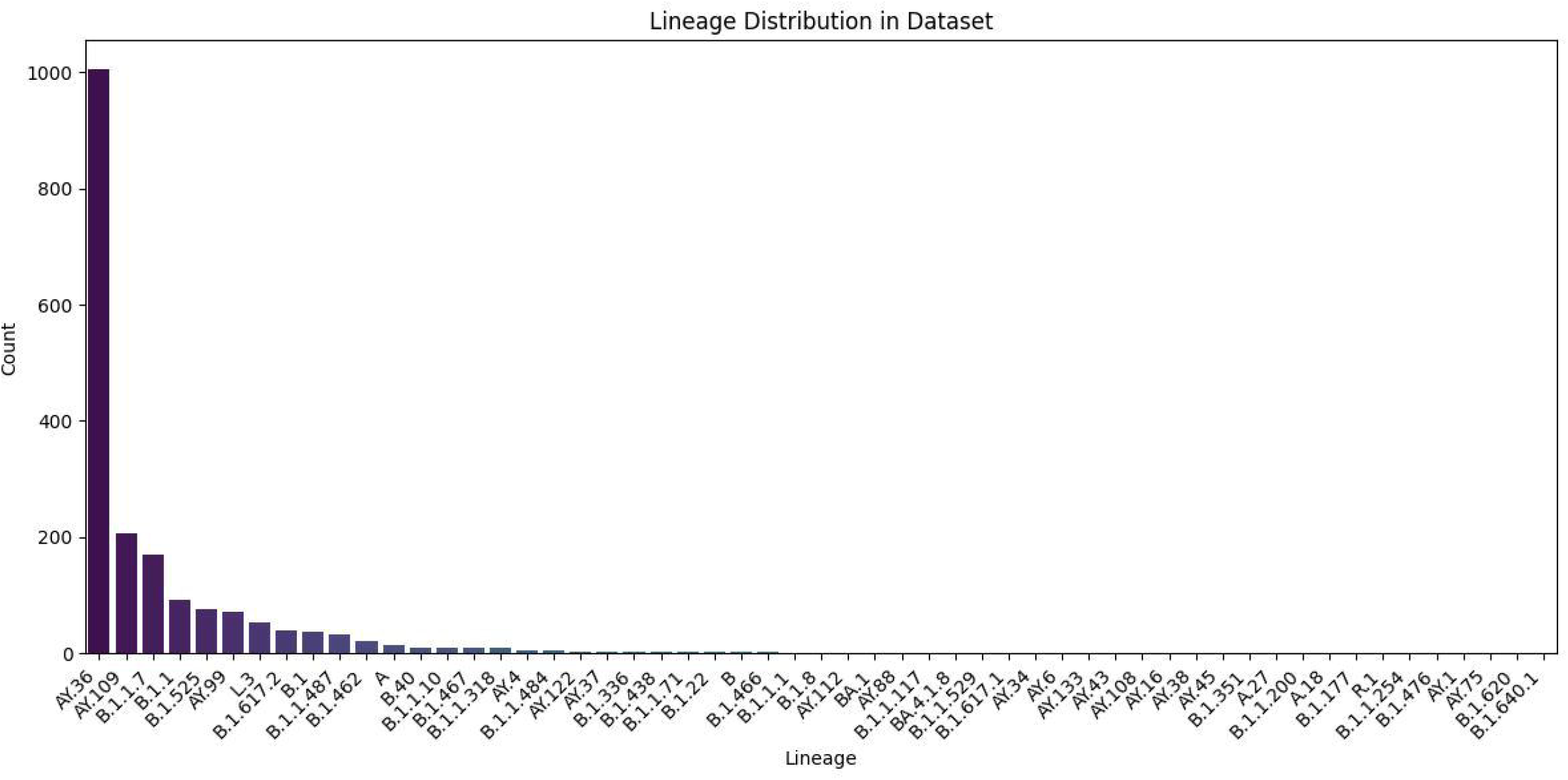

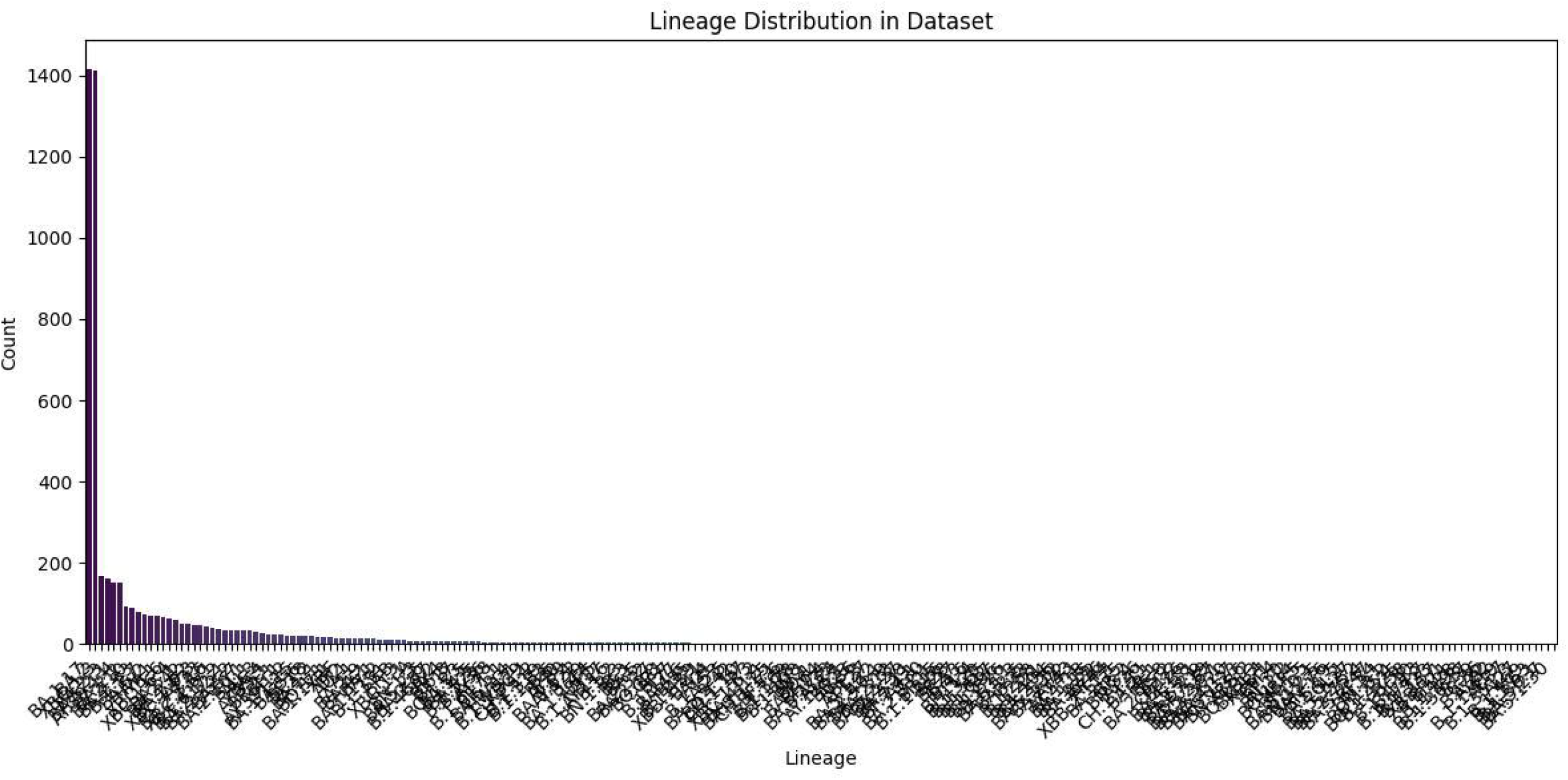

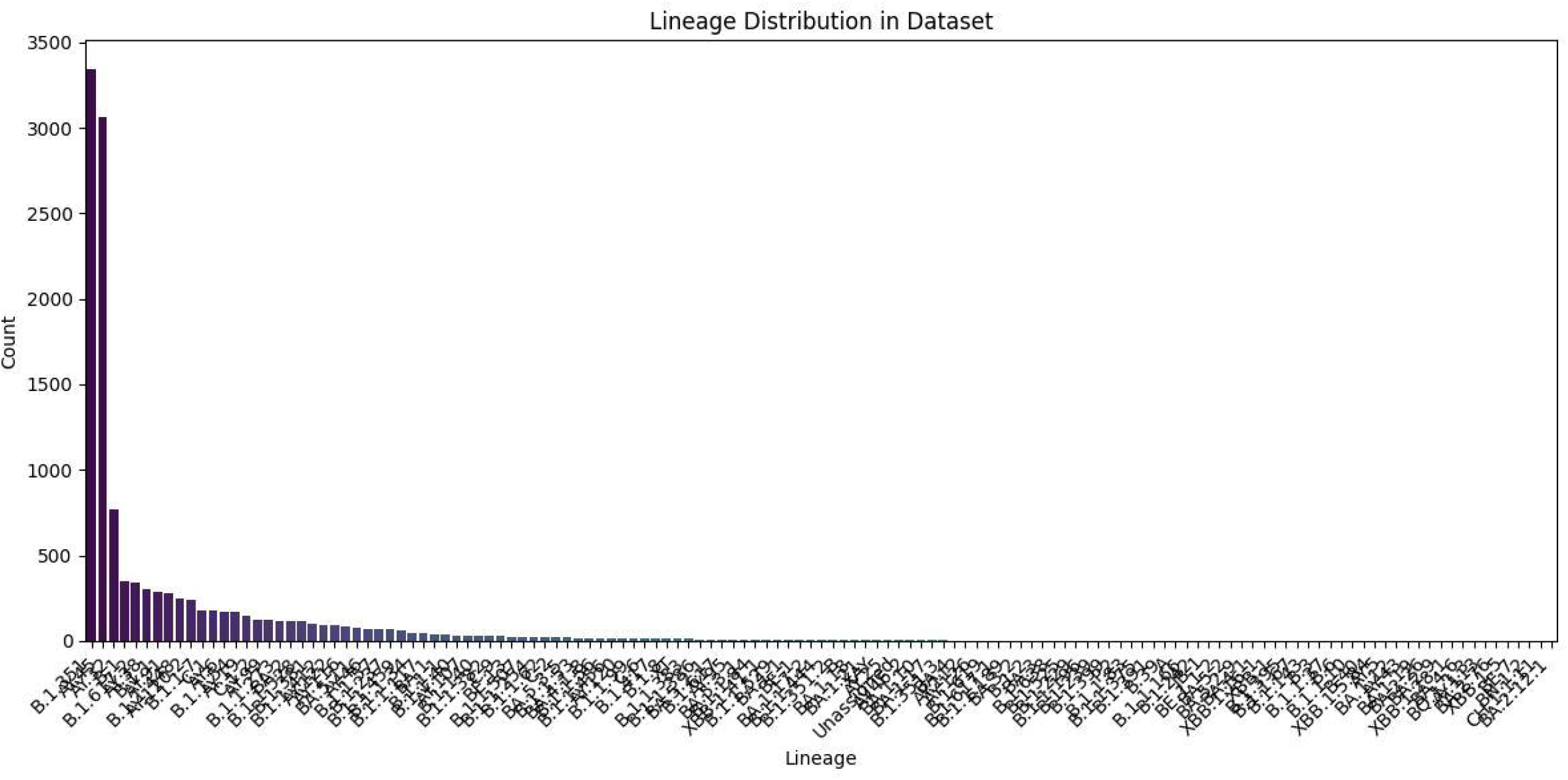

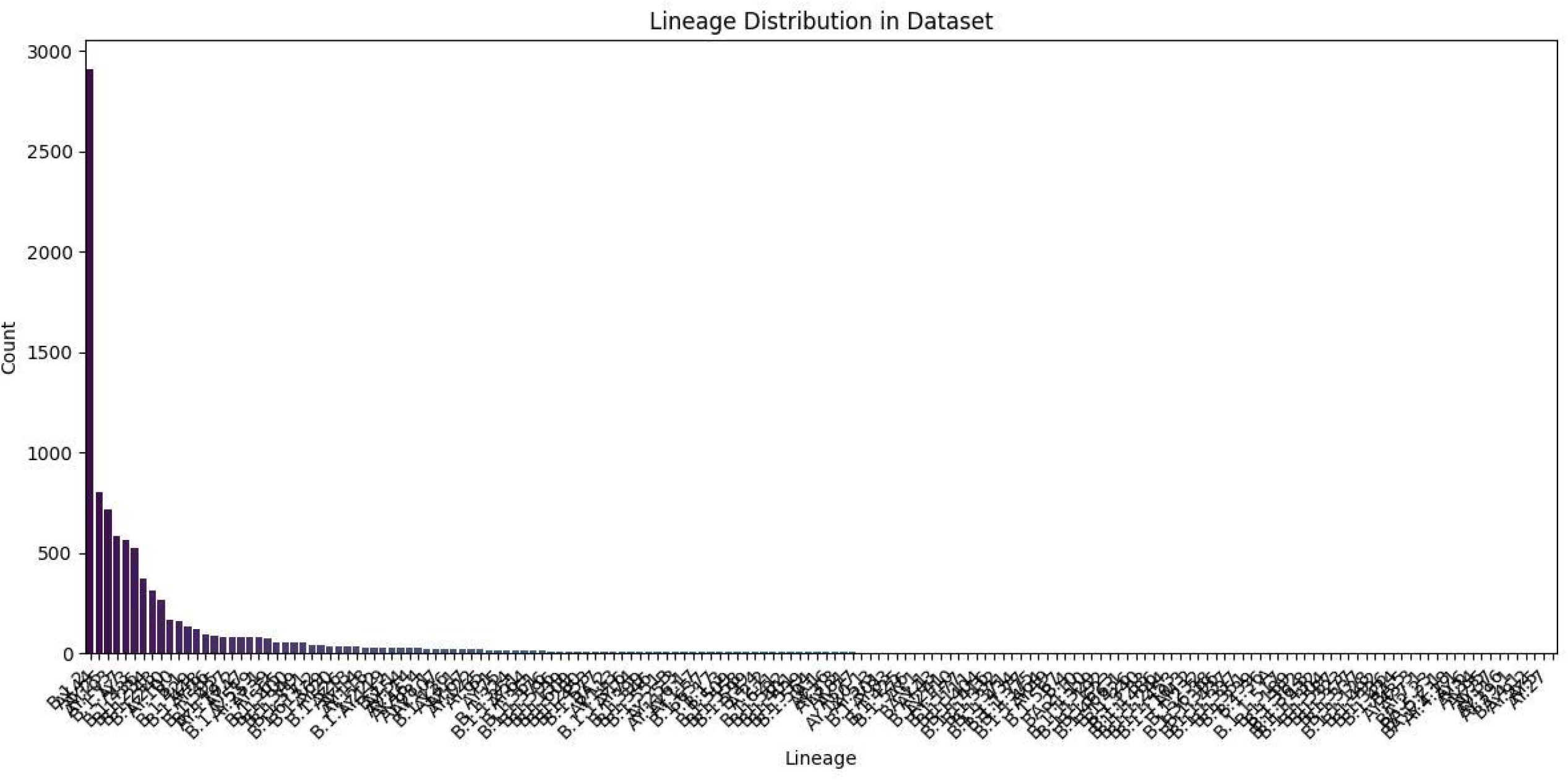

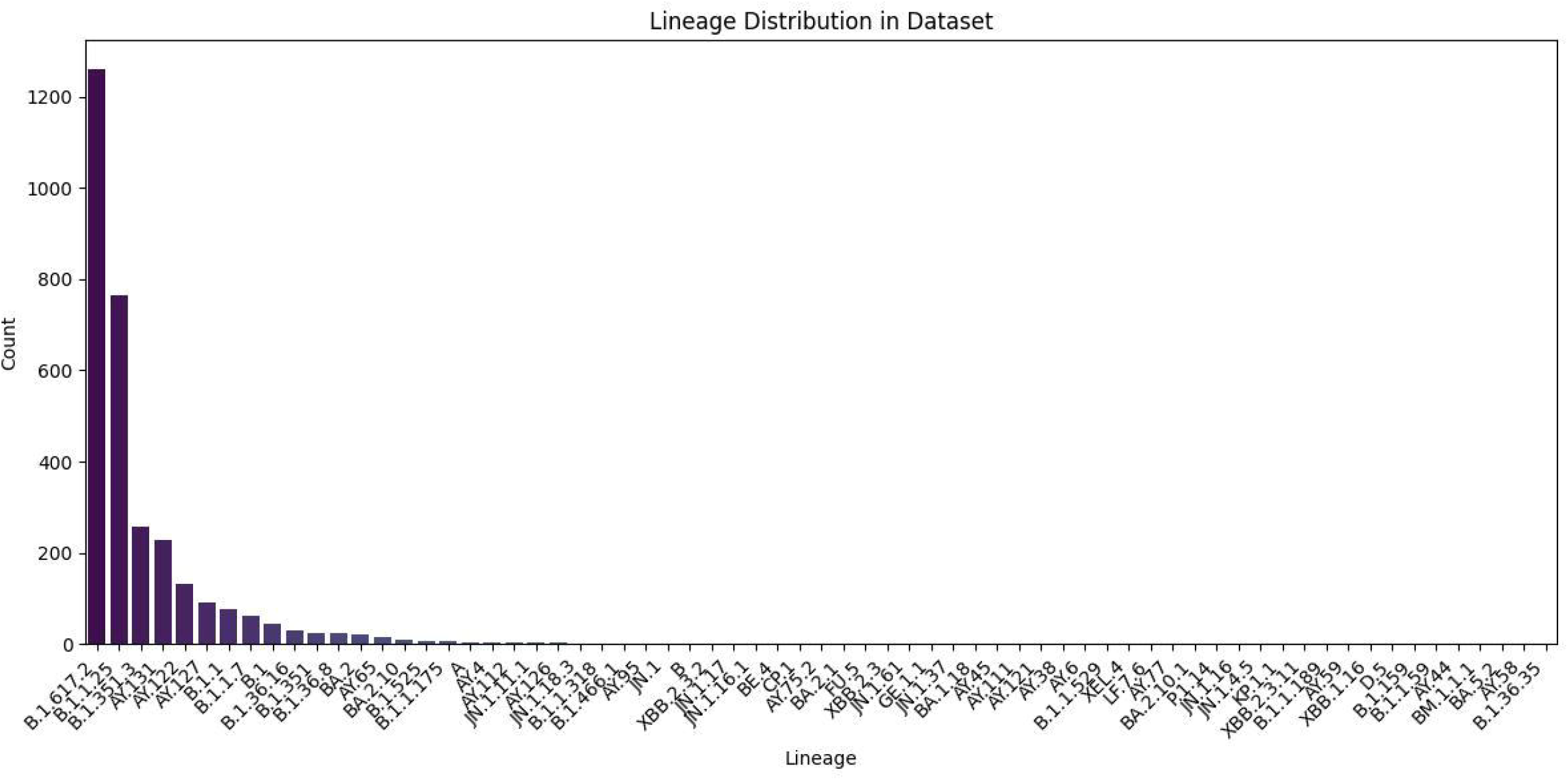

## Footnotes

1 Inputs such as ℳ, *E*, Π, and 𝒟 are derived using the procedures detailed in Sections 2.3 and 2.4.

