## Supplementary material for "Capturing the Mutational Dynamics of SARS-CoV-2 with Graphs"

**Algorithm 1** kNN-Group($G, E, \Pi, k, \mathcal{D}$)

**for all** rows $i$ of $E$ **do**
    mask all but the $k$ highest similarity entries in row $i$ of $S$ (set others to $-\infty$)
    apply the same mask to $E$ to get $E'$ such that $E'_{ij} = E_{ij}$ if $S_{ij}$ is among top-$k$, else $\infty$
    set self-distance $E'_{ii} \leftarrow \infty$
    $d_{\min} \leftarrow \min_j E'_{ij},\ \ s_{\max} \leftarrow \max\{S_{ij} \mid E'_{ij} = d_{\min}\}$
    $N_i \leftarrow \{j \mid E'_{ij} = d_{\min} \wedge S_{ij} = s_{\max}\}$
    $\mathit{common} \leftarrow \bigcap_{v \in \{i\} \cup N_i} M(v)$
    $\mathit{new}$ += Find-Relation($G, \{i\}, N_i, \mathit{common}, \mathcal{D}$)
**end for**
**return** *new*

**Algorithm 2** Find-Relation($G, \mathit{base}, N, \mathit{common}, \mathcal{D}$)

**if** $\mathit{common} = \varnothing$ **then**
    **return** $\varnothing$
**end if**
$P \leftarrow$ mutsetToID(*common*) ▷ find existing node with matching mutation set
**if** $P = \varnothing$ **then**
    $P \leftarrow$ NewID
    store $\mathcal{M}(P) \leftarrow \mathit{common}$ ▷ assign mutation set to new inferred ancestor
    $G$.addNode($P$)
**end if**
**for all** $v \in \{\mathit{base}\} \cup N \setminus \{P\}$ **do**
    **if** $\mathcal{D}(v) \neq \perp$ and $\mathcal{D}(P) \neq \perp$ and $\mathcal{D}(v) < \mathcal{D}(P)$ **then**
        $G$.addEdge($v, P$) ▷ reverse mutation
    **else**
        $G$.addEdge($P, v$) ▷ forward mutation
    **end if**
**end for**
**return** $\{P\}$ if $P$ was newly created, else $\varnothing$

**Algorithm 3** BRIDGE-COMPONENTS$(G, S)$

initialize union–find $UF$ over all node indices
build max-heap $H$ of all off-diagonal $(-S_{ij}, i, j)$ with $S_{ij} > 0$
$new \leftarrow \varnothing$
**while** $UF$ has $> 1$ components **do**
  pop $(\_, i, j)$ with highest similarity from $H$
  **if** $UF.find(i) = UF.find(j)$ **then continue**
  **end if**
  $new$ += FIND_RELATION$(G, \mathrm{id}(i), \mathrm{id}(j), \mathrm{id}(i) \bigcap \mathrm{id}(j), \mathcal{D})$
  $UF$.union$(i, j)$
**end while**
**return** $new$

Table 1: Detailed Pangolin lineage analysis for the inferred variants generated from the Egypt data set.

| **Metric** | **Result** |
|---|---|
| Total variants analyzed | 1058 |
| Assigned lineages | B (all) |
| Conflict values | 0.0 (1024), 0.5 (34) |
| QC status | passed (all) |
| Ambiguity | NaN (unambiguous placements) |
| UShER placement summary | 1022 placed as B(1/1) |
| | 26 placed as B(1/2) B.23(1/2) |
| | 5 placed as B(1/2) B.47(1/2) |
| | 2 placed as B(1/2) B.1.153(1/2) |
| | 2 assigned from designation hash |
| | 1 placed as B(1/2) B.45(1/2) |

Table 2: Detailed Pangolin lineage assignment for the inferred variants generated from the Iran data set.

| **Metric** | **Result** |
|---|---|
| Total variants analyzed | 1373 |
| Assigned lineage | B (all) |
| Conflict | 0.0 (1367), 0.5 (6) |
| QC status | passed (all) |
| Ambiguity | NaN (all) |
| UShER placement summary | 1362 placed as B(1/1) |
| | 4 placed as B(1/2) B.26(1/2) |
| | 2 placed as B(1/2) B.23(1/2) |
| | 5 assigned from the designation hash |

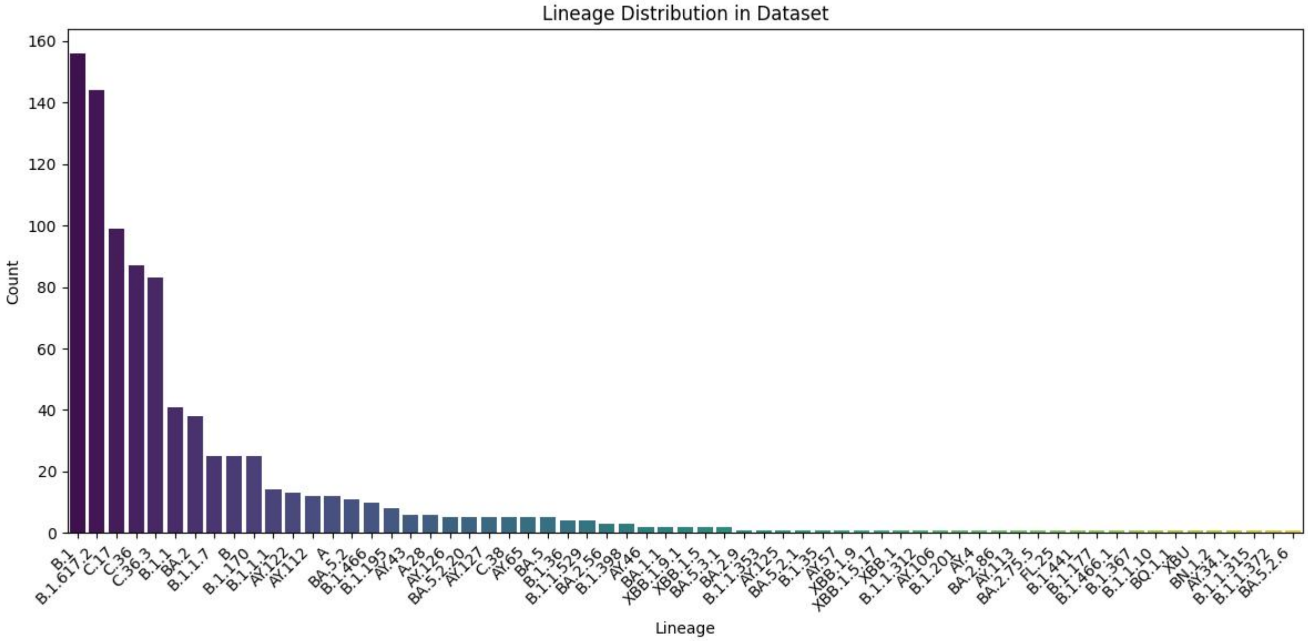


Figure 1: Lineage distribution of the Egypt dataset.

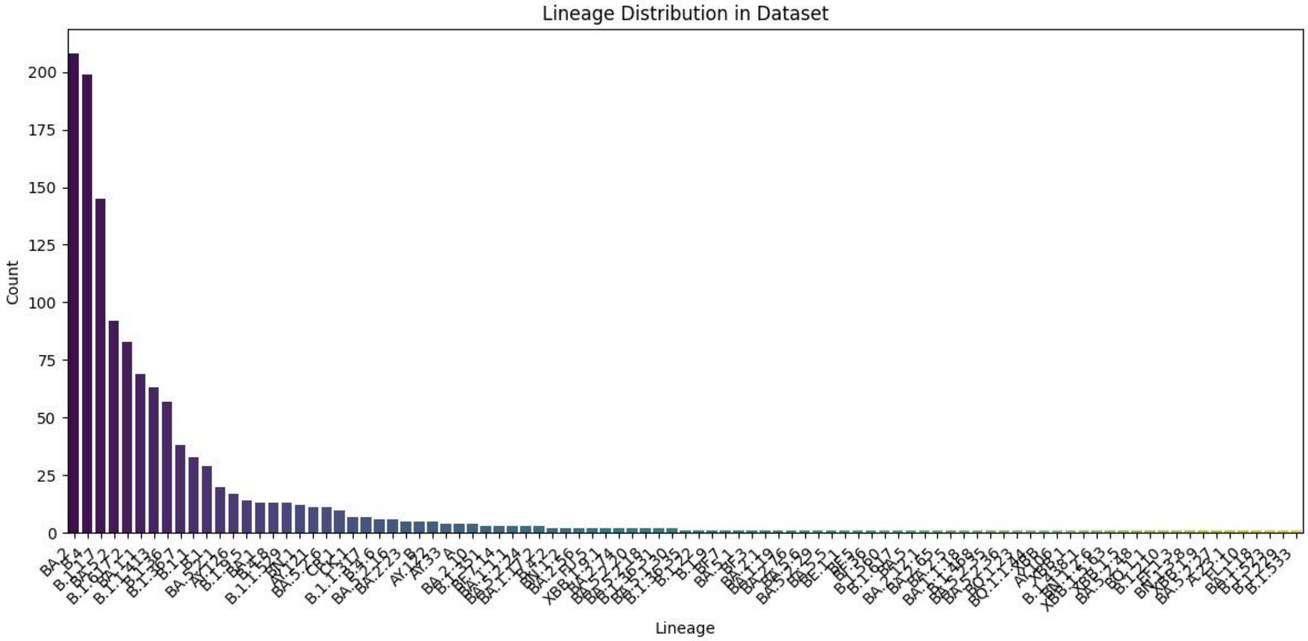


Figure 2: Lineage distribution of the Iran dataset.

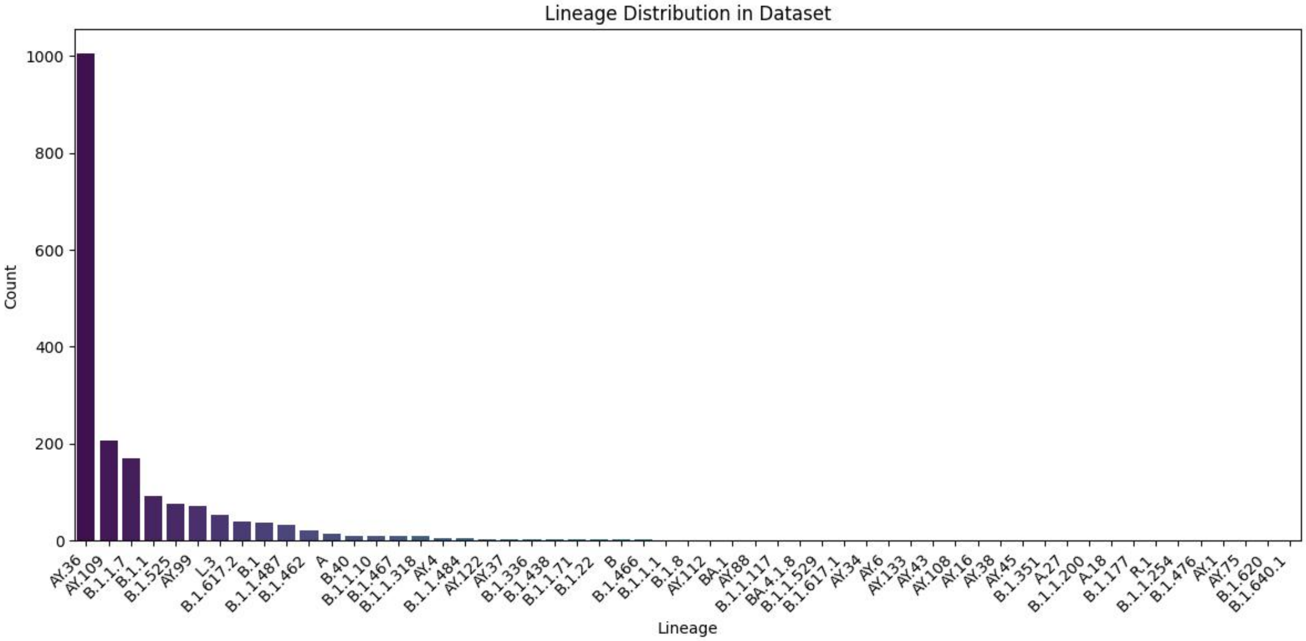


Figure 3: Lineage distribution of the Nigeria dataset.

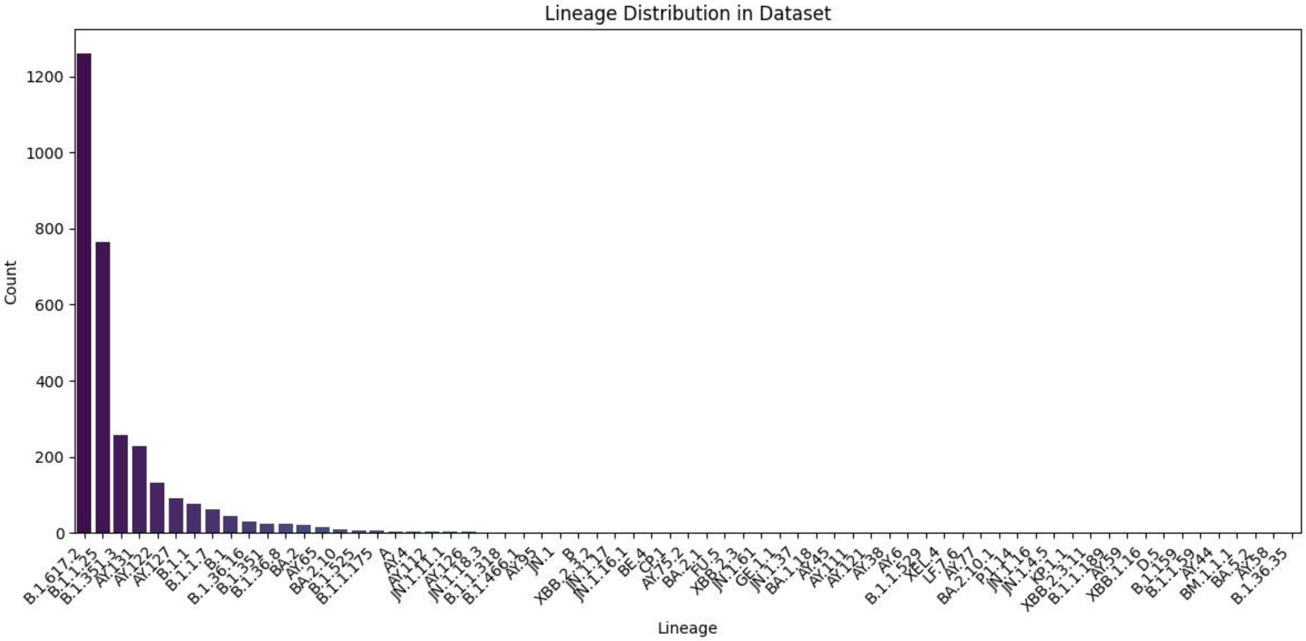


Figure 4: Lineage distribution of the Bangladesh dataset.

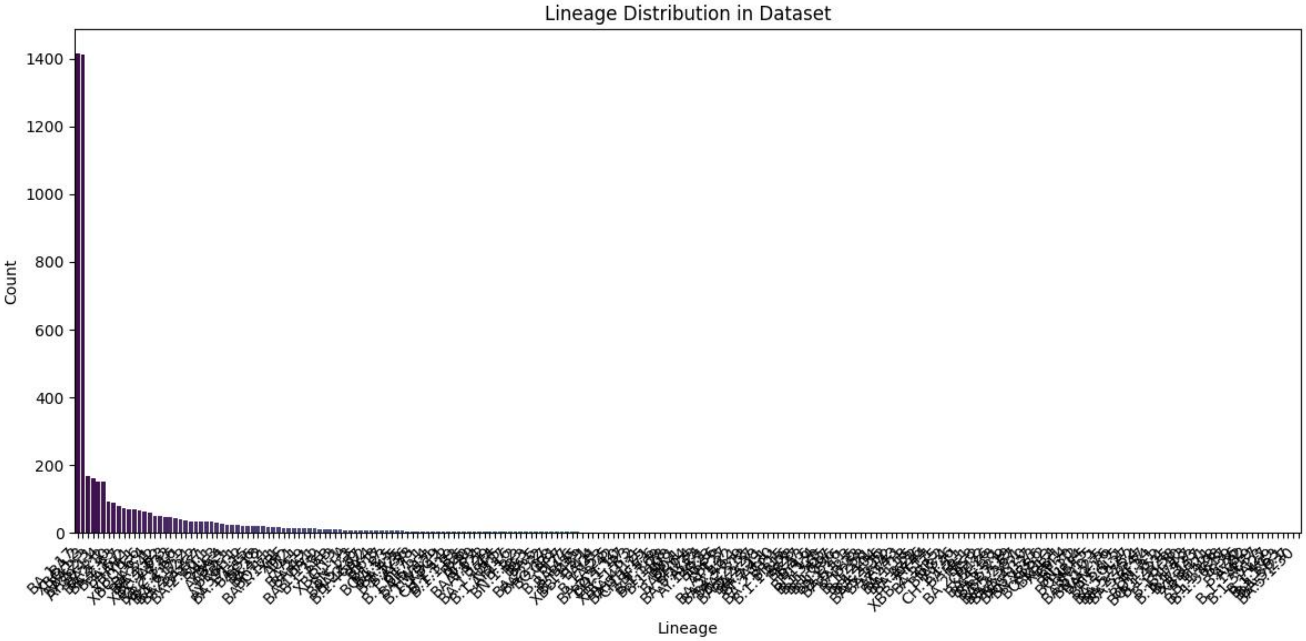


Figure 5: Lineage distribution of the Queensland dataset.

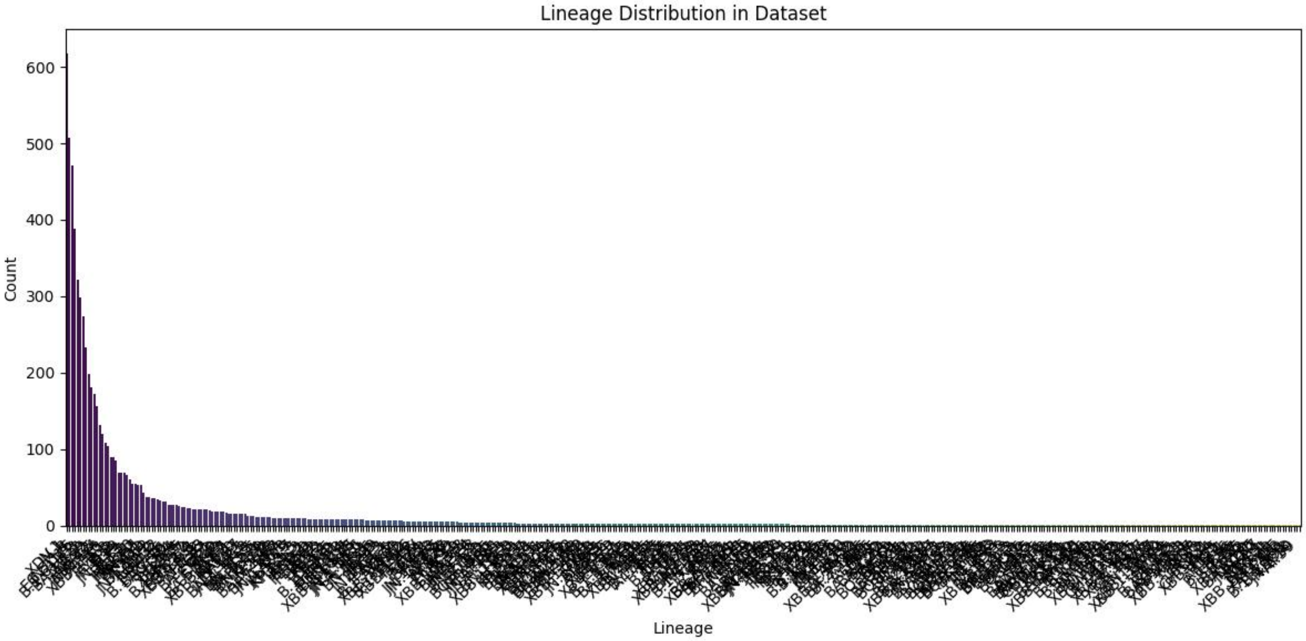


Figure 6: Lineage distribution of the China dataset.

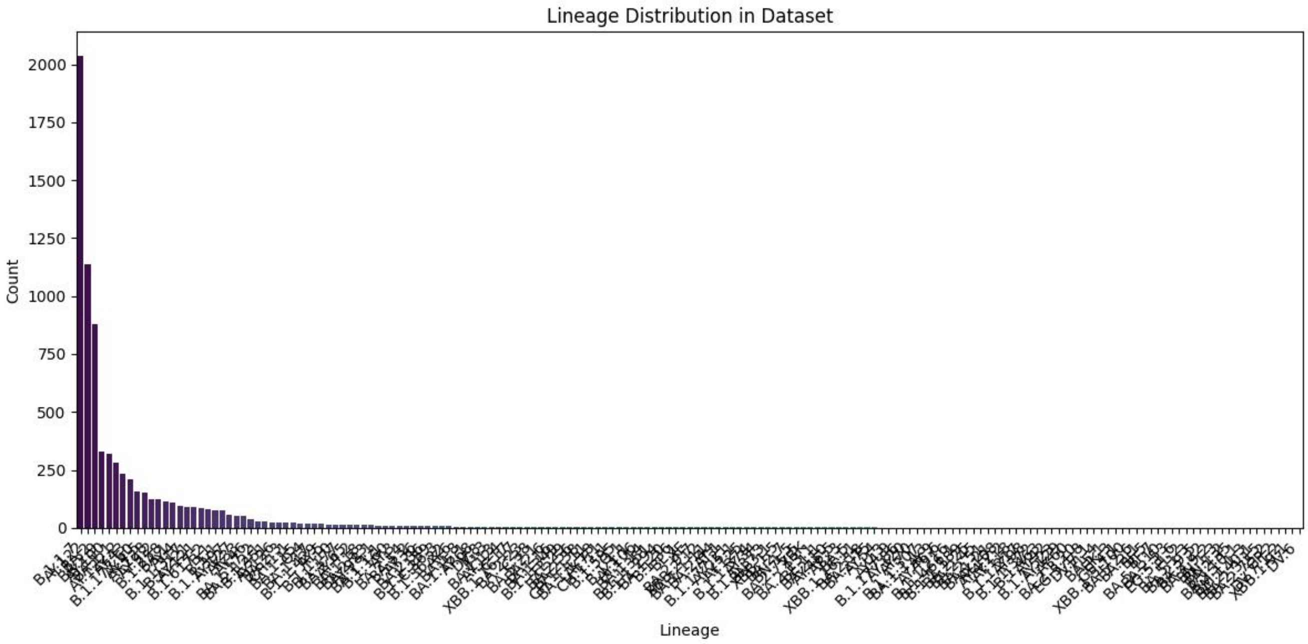


Figure 7: Lineage distribution of the Estonia dataset.

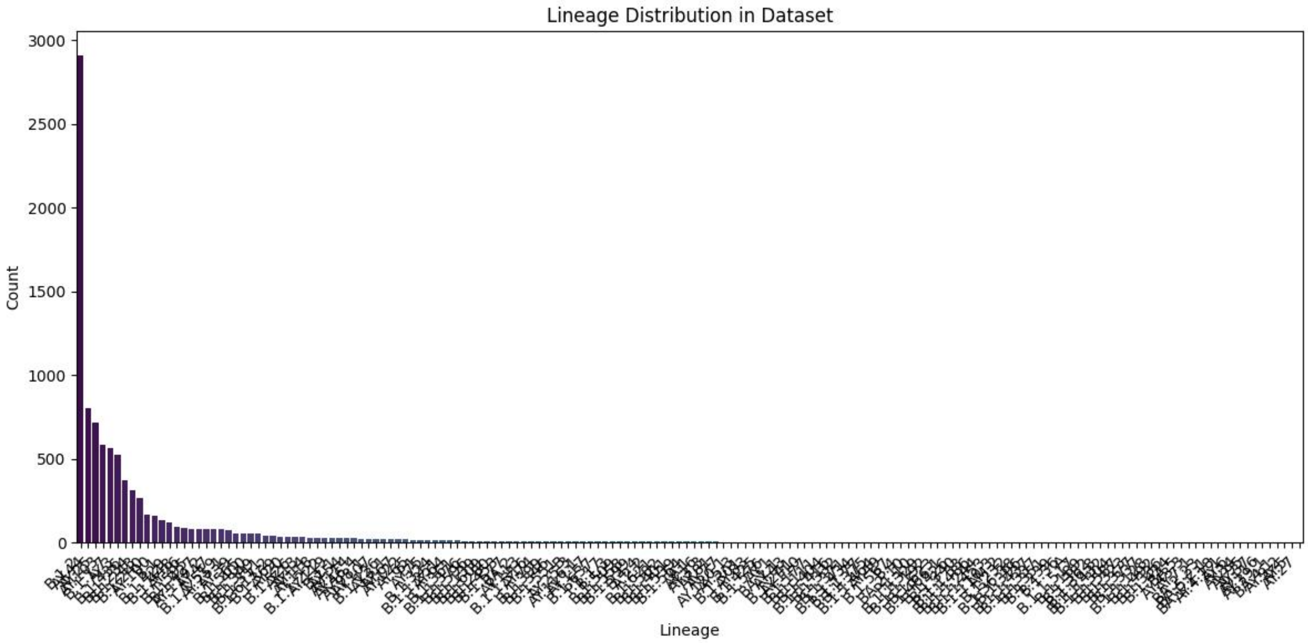


Figure 8: Lineage distribution of the Wyoming dataset.

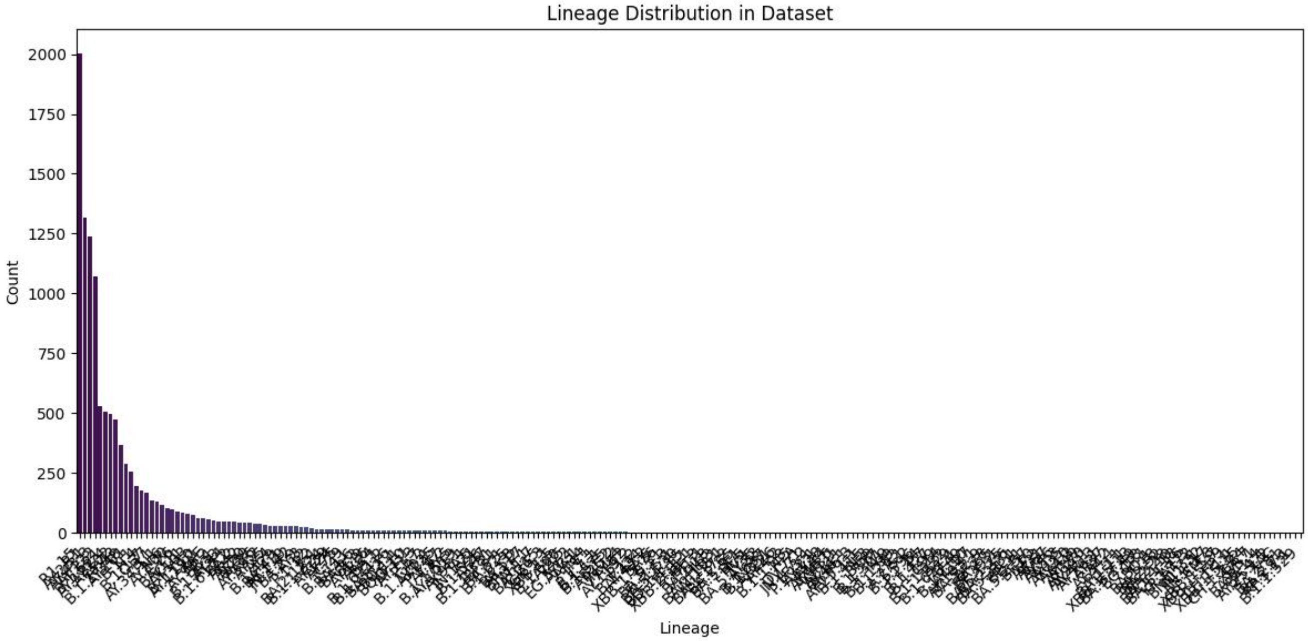


Figure 9: Lineage distribution of the Chile dataset.

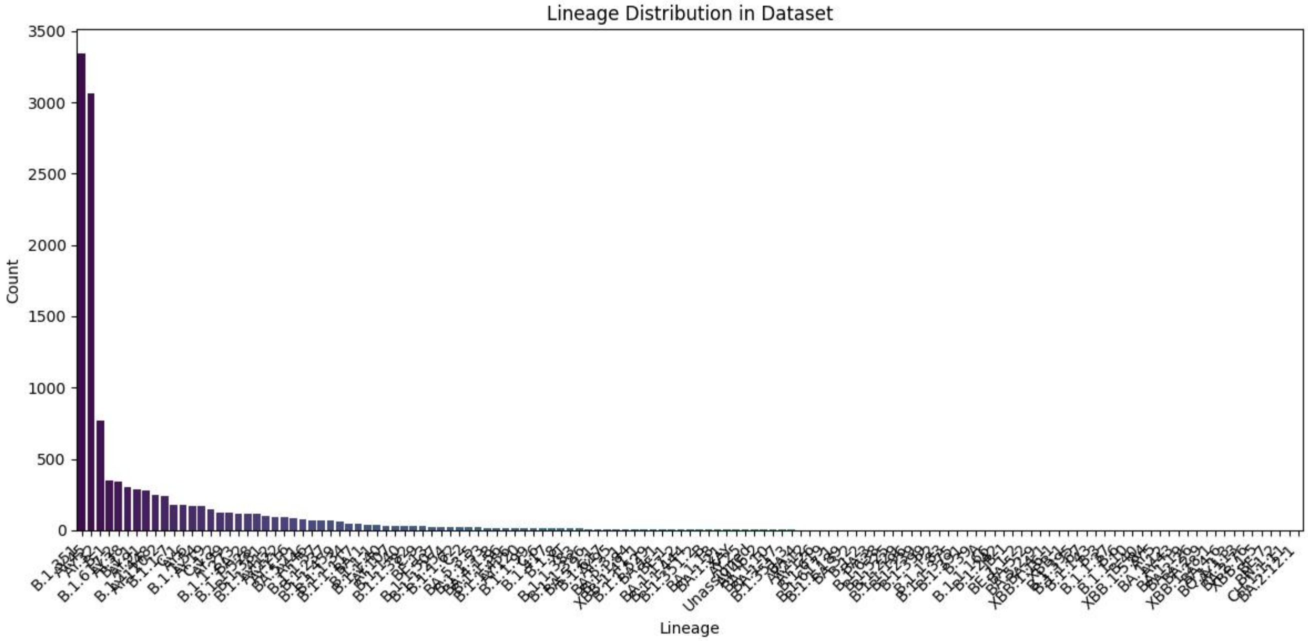


Figure 10: Lineage distribution of the South Africa dataset.

Table 3: Detailed Pangolin lineage assignment for the inferred variants generated from the Nigeria data set.

| **Metric** | **Result** |
|---|---|
| Total variants analyzed | 2013 |
| Assigned lineage | B (all) |
| Conflict | 0.0 (2010), 0.5 (3) |
| QC status | passed (all) |
| Ambiguity | NaN (all) |
| UShER placement summary | 2010 placed as B(1/1) |
| | 3 placed as B(1/2) B.1.14(1/2) |

Table 4: Detailed Pangolin lineage assignment for the inferred variants generated from the Bangladesh data set.

| **Metric** | **Result** |
|---|---|
| Total variants analyzed | 1373 |
| Assigned lineage | B (3582), B.19 (7) |
| Conflict | 0.0 (3547), 0.5 (42) |
| QC status | passed (all) |
| Ambiguity | NaN (all) |
| UShER placement summary | 3540 placed as B(1/1) |
| | 13 placed as B(1/2) B.26(1/2) |
| | 11 placed as B(1/2) B.12(1/2) |
| | 7 placed as B.19(1/1) |
| | 5 placed as B(1/2) B.23(1/2) |
| | 5 placed as B(1/2) B.47(1/2) |
| | 3 placed as B(1/2) B.11(1/2) |
| | 3 placed as B(1/2) B.1.1(1/2) |
| | 2 placed as B(1/2) B.1.14(1/2) |

Table 5: Detailed Pangolin lineage assignment for the inferred variants generated from the Queensland data set.

| **Metric** | **Result** |
|---|---|
| Total variants analyzed | 5582 |
| Assigned lineage | B (all) |
| Conflict | 0.0 (5539), 0.5 (36), 0.67 (7) |
| QC status | passed (all) |
| Ambiguity | NaN (all) |
| UShER placement summary | 5538 placed as B(1/1) |
| | 19 placed as B(1/2) B.26(1/2) |
| | 10 placed as B(1/2) B.1.14(1/2) |
| | 7 placed as B(1/3) B.1.14(1/3) B.47(1/3) |
| | 3 placed as B(1/2) B.19(1/2) |
| | 2 placed as B(1/2) B.47(1/2) |
| | 2 placed as B(1/2) B.4(1/2) |
| | 1 assigned from the designation hash |

Table 6: Detailed Pangolin lineage assignment for the inferred variants generated from the China data set.

| **Metric** | **Result** |
|---|---|
| Total variants analyzed | 6066 |
| Assigned lineage | B (all) |
| Conflict | 0.0 (6050), 0.5 (13), 0.67 (3) |
| QC status | passed (all) |
| Ambiguity | NaN (all) |
| UShER placement summary | 6002 placed as B(1/1) |
| | 14 assigned from designation hash |
| | 4 placed as B(1/2) B.4(1/2) |
| | 3 placed as B(1/2) B.1.14(1/2) |
| | 3 placed as B(1/2) B.11(1/2) |
| | 2 placed as B(1/3) B.1.14(1/3) B.47(1/3) |
| | 2 placed as B(1/2) B.23(1/2) |
| | 1 placed as B(1/3) B.12(1/3) B.23(1/3) |
| | 1 placed as B(1/2) B.12(1/2) |

Table 7: Detailed Pangolin lineage assignment for the inferred variants generated from the Estonia data set.

| **Metric** | **Result** |
|---|---|
| Total variants analyzed | 5524 |
| Assigned lineage | B (all) |
| Conflict | 0.0 (5451), 0.5 (38), 0.67 (35) |
| QC status | passed (all) |
| Ambiguity | NaN (all) |
| UShER placement summary | 5439 placed as B(1/1) |
| | 35 placed as B(1/3) B.1.14(1/3) B.26(1/3) |
| | 14 placed as B(1/2) B.26(1/2) |
| | 12 placed as B(2/2) |
| | 10 placed as B(1/2) B.1.14(1/2) |
| | 9 placed as B(1/2) B.11(1/2) |
| | 3 placed as B(1/2) B.23(1/2) |
| | 1 placed as B(1/2) B.12(1/2) |
| | 1 placed as B(1/2) B.47(1/2) |

Table 8: Detailed Pangolin lineage assignment for the inferred variants generated from the Wyoming data set.

| **Metric** | **Result** |
|---|---|
| Total variants analyzed | 8523 |
| Assigned lineage | B (8514), A (7), B.19 (2) |
| Conflict | 0.0 (8261), 0.5 (262) |
| QC status | passed (all) |
| Ambiguity | NaN (all) |
| UShER placement summary | 8252 placed as B(1/1) |
| | 183 placed as B(1/2) B.1.14(1/2) |
| | 38 placed as B(1/2) B.23(1/2) |
| | 24 placed as B(1/2) B.26(1/2) |
| | 7 placed as A(1/1) |
| | 7 placed as B(1/2) B.11(1/2) |
| | 6 placed as B(1/2) B.47(1/2) |
| | 3 placed as B(1/2) B.4(1/2) |
| | 2 placed as B.19(1/1) |
| | 1 placed as B(1/2) B.55(1/2) |

Table 9: Detailed Pangolin lineage assignment for the inferred variants generated from the Chile data set.

| **Metric** | **Result** |
|---|---|
| Total variants analyzed | 12251 |
| Assigned lineage | B (12249), B.19 (2) |
| Conflict | 0.0 (12180), 0.5 (71) |
| QC status | passed (all) |
| Ambiguity | NaN (all) |
| UShER placement summary | 12166 placed as B(1/1) |
| | 31 placed as B(1/2) B.26(1/2) |
| | 17 placed as B(1/2) B.47(1/2) |
| | 14 placed as B(1/2) B.11(1/2) |
| | 12 placed as B(2/2) |
| | 6 placed as B(1/2) B.1.14(1/2) |
| | 3 placed as B(1/2) B.23(1/2) |
| | 2 placed as B.19(1/1) |

Table 10: Detailed Pangolin lineage assignment for the inferred variants generated from the South Africa data set.

| **Metric** | **Result** |
|---|---|
| Total variants analyzed | 12912 |
| Assigned lineage | B (12911), A (1) |
| Conflict | 0.0 (12869), 0.5 (43) |
| QC status | passed (all) |
| Ambiguity | NaN (all) |
| UShER placement summary | 12864 placed as B(1/1) |
| | 21 placed as B(1/2) B.26(1/2) |
| | 5 placed as B(1/2) B.23(1/2) |
| | 4 placed as B(1/2) B.11(1/2) |
| | 4 placed as B(2/2) |
| | 3 placed as B(1/2) B.4(1/2) |
| | 3 placed as B(1/2) B.19(1/2) |
| | 3 placed as B(1/2) B.1.14(1/2) |
| | 2 placed as B(1/2) B.56(1/2) |
| | 1 placed as B(1/2) B.47(1/2) |
| | 1 placed as A(1/1) |
| | 1 placed as B(1/2) B.55(1/2) |

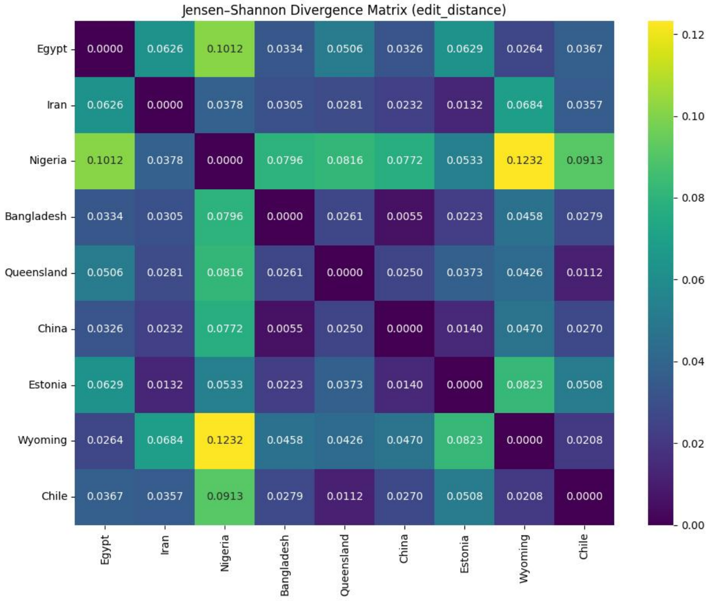


Figure 11: Jensen–Shannon divergence matrix for *edit distance* across ten regional MLGs. Higher values indicate greater dissimilarity in mutational connectivity patterns.

Table 11: Experiment results for the ten data sets on the node classification task

| Data Set | Metric | GCN | GraphSAGE | GAT | GGNN | MLP |
|---|---|---|---|---|---|---|
| **Egypt** | Acc | 0.8713 ± 0.0141 | 0.7937 ± 0.0228 | 0.6514 ± 0.0603 | 0.7826 ± 0.0637 | 0.9625 ± 0.0208 |
| | F1 | 0.8766 ± 0.0126 | 0.8039 ± 0.0256 | 0.6659 ± 0.0516 | 0.7846 ± 0.0726 | 0.9808 ± 0.0109 |
| | AUROC | 0.9981 ± 0.0004 | nan ± nan | nan ± nan | nan ± nan | nan ± nan |
| | AUPRC | 0.9612 ± 0.0093 | 0.6050 ± 0.0266 | 0.5263 ± 0.0379 | 0.5959 ± 0.0365 | 0.0333 ± 0.0000 |
| **Iran** | Acc | 0.8079 ± 0.0263 | 0.8749 ± 0.0223 | 0.6946 ± 0.0283 | 0.8828 ± 0.0146 | 0.9685 ± 0.0103 |
| | F1 | 0.8251 ± 0.0213 | 0.8732 ± 0.0225 | 0.7145 ± 0.0262 | 0.8818 ± 0.0165 | 0.9840 ± 0.0053 |
| | AUROC | 0.9971 ± 0.0006 | nan ± nan | nan ± nan | nan ± nan | nan ± nan |
| | AUPRC | 0.9289 ± 0.0129 | 0.6270 ± 0.0351 | 0.5593 ± 0.0329 | 0.6472 ± 0.0402 | 0.0270 ± 0.0000 |
| **Nigeria** | Acc | 0.8387 ± 0.0126 | 0.9416 ± 0.0126 | 0.7612 ± 0.0302 | 0.9511 ± 0.0149 | 0.9624 ± 0.0087 |
| | F1 | 0.8529 ± 0.0116 | 0.9453 ± 0.0109 | 0.7823 ± 0.0248 | 0.9524 ± 0.0132 | 0.9808 ± 0.0045 |
| | AUROC | 0.9947 ± 0.0012 | nan ± nan | nan ± nan | nan ± nan | nan ± nan |
| | AUPRC | 0.9237 ± 0.0100 | 0.7377 ± 0.0402 | 0.6712 ± 0.0444 | 0.7388 ± 0.0435 | 0.0370 ± 0.0000 |
| **Bangladesh** | Acc | 0.8183 ± 0.0141 | 0.8758 ± 0.0180 | 0.7267 ± 0.0903 | 0.9154 ± 0.0484 | 0.9833 ± 0.0094 |
| | F1 | 0.8417 ± 0.0110 | 0.9034 ± 0.0124 | 0.7556 ± 0.0754 | 0.9311 ± 0.0343 | 0.9916 ± 0.0048 |
| | AUROC | 0.9936 ± 0.0009 | nan ± nan | nan ± nan | nan ± nan | nan ± nan |
| | AUPRC | 0.9178 ± 0.0072 | 0.6231 ± 0.0648 | 0.5831 ± 0.0717 | 0.6592 ± 0.0742 | 0.0435 ± 0.0000 |
| **Queensland (Australia)** | Acc | 0.5392 ± 0.0173 | 0.7593 ± 0.0157 | 0.4702 ± 0.0273 | 0.7861 ± 0.0291 | 0.8512 ± 0.0239 |
| | F1 | 0.5805 ± 0.0189 | 0.7883 ± 0.0140 | 0.5015 ± 0.0320 | 0.8101 ± 0.0230 | 0.9195 ± 0.0141 |
| | AUROC | 0.9943 ± 0.0003 | nan ± nan | nan ± nan | nan ± nan | nan ± nan |
| | AUPRC | 0.8165 ± 0.0068 | 0.6475 ± 0.0137 | 0.5047 ± 0.0240 | 0.6792 ± 0.0164 | 0.0101 ± 0.0000 |
| **China** | Acc | 0.4737 ± 0.0186 | 0.5744 ± 0.0358 | 0.2811 ± 0.0382 | 0.5906 ± 0.0447 | 0.5918 ± 0.0496 |
| | F1 | 0.4925 ± 0.0196 | 0.6363 ± 0.0339 | 0.3139 ± 0.0421 | 0.6451 ± 0.0466 | 0.7195 ± 0.0376 |
| | AUROC | 0.9949 ± 0.0003 | nan ± nan | nan ± nan | nan ± nan | nan ± nan |
| | AUPRC | 0.6997 ± 0.0078 | 0.2301 ± 0.0128 | 0.1578 ± 0.0113 | 0.2276 ± 0.0138 | 0.0463 ± 0.0072 |
| **Estonia** | Acc | 0.5499 ± 0.0229 | 0.9197 ± 0.0076 | 0.4706 ± 0.0310 | 0.9353 ± 0.0072 | 0.8816 ± 0.0258 |
| | F1 | 0.5869 ± 0.0227 | 0.9274 ± 0.0062 | 0.5053 ± 0.0371 | 0.9418 ± 0.0056 | 0.9185 ± 0.0191 |
| | AUROC | 0.9899 ± 0.0006 | nan ± nan | nan ± nan | nan ± nan | nan ± nan |
| | AUPRC | 0.7213 ± 0.0095 | 0.4258 ± 0.0110 | 0.2827 ± 0.0072 | 0.4505 ± 0.0091 | 0.2736 ± 0.0095 |
| **Wyoming (USA)** | Acc | 0.5677 ± 0.0238 | 0.9020 ± 0.0120 | 0.4967 ± 0.0435 | 0.9124 ± 0.0268 | 0.8987 ± 0.0182 |
| | F1 | 0.6002 ± 0.0245 | 0.9187 ± 0.0082 | 0.5559 ± 0.0426 | 0.9243 ± 0.0220 | 0.9409 ± 0.0111 |
| | AUROC | 0.9883 ± 0.0007 | nan ± nan | nan ± nan | nan ± nan | nan ± nan |
| | AUPRC | 0.7020 ± 0.0094 | 0.6373 ± 0.0199 | 0.4582 ± 0.0257 | 0.6552 ± 0.0209 | 0.2561 ± 0.0212 |
| **Chile** | Acc | 0.5970 ± 0.0195 | 0.8496 ± 0.0299 | 0.4893 ± 0.0462 | 0.8676 ± 0.0319 | 0.7450 ± 0.0312 |
| | F1 | 0.6234 ± 0.0169 | 0.8690 ± 0.0199 | 0.5455 ± 0.0489 | 0.8841 ± 0.0223 | 0.8535 ± 0.0208 |
| | AUROC | 0.9897 ± 0.0003 | nan ± nan | nan ± nan | nan ± nan | nan ± nan |
| | AUPRC | 0.6982 ± 0.0086 | 0.2931 ± 0.0105 | 0.1844 ± 0.0094 | 0.3130 ± 0.0170 | 0.0087 ± 0.0000 |
| **South Africa** | Acc | 0.6398 ± 0.0158 | 0.8553 ± 0.0121 | 0.5968 ± 0.0253 | 0.8691 ± 0.0148 | 0.9178 ± 0.0197 |
| | F1 | 0.6719 ± 0.0123 | 0.8654 ± 0.0090 | 0.6323 ± 0.0176 | 0.8749 ± 0.0151 | 0.9570 ± 0.0109 |
| | AUROC | 0.9836 ± 0.0015 | nan ± nan | nan ± nan | nan ± nan | nan ± nan |
| | AUPRC | 0.7475 ± 0.0056 | 0.5226 ± 0.0100 | 0.3679 ± 0.0086 | 0.5481 ± 0.0153 | 0.0128 ± 0.0000 |

Table 12: Experiment results for the ten data sets on the link prediction task

| Data Set | Metric | GraphSAGE | GAT | VGAE | GGNN | MLP |
|---|---|---|---|---|---|---|
| **Egypt** | Acc | 0.9512 ± 0.0222 | 0.6937 ± 0.0548 | 0.9117 ± 0.0574 | 0.9852 ± 0.0038 | 0.6936 ± 0.0327 |
| | F1 | 0.9508 ± 0.0217 | 0.7323 ± 0.0466 | 0.9079 ± 0.0646 | 0.6477 ± 0.0829 | 0.6919 ± 0.0326 |
| | AUROC | 0.9717 ± 0.0165 | 0.8238 ± 0.0487 | 0.9634 ± 0.0229 | 0.9890 ± 0.0077 | 0.7609 ± 0.0350 |
| | AUPRC | 0.9461 ± 0.0299 | 0.8106 ± 0.0575 | 0.9473 ± 0.0319 | 0.7897 ± 0.0637 | 0.7719 ± 0.0509 |
| **Iran** | Acc | 0.9560 ± 0.0146 | 0.7027 ± 0.0354 | 0.9347 ± 0.0168 | 0.9950 ± 0.0034 | 0.8014 ± 0.0397 |
| | F1 | 0.9584 ± 0.0121 | 0.7560 ± 0.0271 | 0.9384 ± 0.0162 | 0.2305 ± 0.3266 | 0.8011 ± 0.0398 |
| | AUROC | 0.9663 ± 0.0202 | 0.8072 ± 0.0380 | 0.9631 ± 0.0211 | 0.9551 ± 0.0566 | 0.8739 ± 0.0350 |
| | AUPRC | 0.9428 ± 0.0335 | 0.7799 ± 0.0290 | 0.9372 ± 0.0412 | 0.3777 ± 0.2490 | 0.8659 ± 0.0382 |
| **Nigeria** | Acc | 0.9502 ± 0.0088 | 0.6144 ± 0.0369 | 0.9502 ± 0.0150 | 0.9865 ± 0.0034 | 0.6608 ± 0.0355 |
| | F1 | 0.9526 ± 0.0079 | 0.6833 ± 0.0229 | 0.9530 ± 0.0138 | 0.2879 ± 0.1015 | 0.6598 ± 0.0357 |
| | AUROC | 0.9710 ± 0.0122 | 0.7244 ± 0.0333 | 0.9699 ± 0.0143 | 0.9616 ± 0.0282 | 0.7204 ± 0.0291 |
| | AUPRC | 0.9494 ± 0.0286 | 0.7124 ± 0.0390 | 0.9455 ± 0.0261 | 0.3010 ± 0.1022 | 0.7349 ± 0.0390 |
| **Bangladesh** | Acc | 0.9639 ± 0.0130 | 0.6128 ± 0.0329 | 0.9542 ± 0.0160 | 0.9907 ± 0.0024 | 0.6977 ± 0.0268 |
| | F1 | 0.9637 ± 0.0133 | 0.6862 ± 0.0245 | 0.9532 ± 0.0163 | 0.3295 ± 0.1230 | 0.6966 ± 0.0265 |
| | AUROC | 0.9764 ± 0.0110 | 0.7438 ± 0.0179 | 0.9782 ± 0.0153 | 0.9855 ± 0.0053 | 0.7746 ± 0.0280 |
| | AUPRC | 0.9533 ± 0.0243 | 0.7784 ± 0.0194 | 0.9625 ± 0.0312 | 0.3776 ± 0.1059 | 0.7768 ± 0.0292 |
| **Queensland (Australia)** | Acc | 0.9478 ± 0.0064 | 0.6184 ± 0.0553 | 0.9402 ± 0.0145 | 0.9853 ± 0.0019 | 0.8163 ± 0.0131 |
| | F1 | 0.9495 ± 0.0064 | 0.7172 ± 0.0286 | 0.9431 ± 0.0127 | 0.3861 ± 0.1428 | 0.8158 ± 0.0134 |
| | AUROC | 0.9685 ± 0.0072 | 0.8280 ± 0.0120 | 0.9684 ± 0.0091 | 0.9699 ± 0.0175 | 0.8853 ± 0.0109 |
| | AUPRC | 0.9436 ± 0.0176 | 0.8105 ± 0.0248 | 0.9522 ± 0.0144 | 0.5006 ± 0.0699 | 0.8603 ± 0.0173 |
| **China** | Acc | 0.9615 ± 0.0048 | 0.7052 ± 0.0459 | 0.9492 ± 0.0137 | 0.9872 ± 0.0025 | 0.8730 ± 0.0132 |
| | F1 | 0.9619 ± 0.0046 | 0.7647 ± 0.0277 | 0.9514 ± 0.0125 | 0.5543 ± 0.1926 | 0.8730 ± 0.0132 |
| | AUROC | 0.9791 ± 0.0044 | 0.8691 ± 0.0095 | 0.9757 ± 0.0077 | 0.9832 ± 0.0080 | 0.9426 ± 0.0117 |
| | AUPRC | 0.9623 ± 0.0087 | 0.8523 ± 0.0145 | 0.9645 ± 0.0129 | 0.7139 ± 0.0357 | 0.9328 ± 0.0164 |
| **Estonia** | Acc | 0.9463 ± 0.0095 | 0.5622 ± 0.0567 | 0.9402 ± 0.0151 | 0.9763 ± 0.0035 | 0.8267 ± 0.0084 |
| | F1 | 0.9481 ± 0.0095 | 0.6960 ± 0.0283 | 0.9424 ± 0.0148 | 0.3618 ± 0.1886 | 0.8258 ± 0.0084 |
| | AUROC | 0.9741 ± 0.0075 | 0.7786 ± 0.0339 | 0.9735 ± 0.0094 | 0.9678 ± 0.0056 | 0.8905 ± 0.0072 |
| | AUPRC | 0.9572 ± 0.0156 | 0.7545 ± 0.0237 | 0.9610 ± 0.0148 | 0.5385 ± 0.0583 | 0.8544 ± 0.0099 |
| **Wyoming (USA)** | Acc | 0.9335 ± 0.0078 | 0.5279 ± 0.0155 | 0.9234 ± 0.0108 | 0.9792 ± 0.0035 | 0.6501 ± 0.0210 |
| | F1 | 0.9352 ± 0.0075 | 0.6704 ± 0.0048 | 0.9271 ± 0.0093 | 0.4081 ± 0.0795 | 0.6450 ± 0.0298 |
| | AUROC | 0.9600 ± 0.0066 | 0.6756 ± 0.0139 | 0.9546 ± 0.0084 | 0.9750 ± 0.0035 | 0.6987 ± 0.0205 |
| | AUPRC | 0.9287 ± 0.0143 | 0.6437 ± 0.0177 | 0.9287 ± 0.0132 | 0.4885 ± 0.0619 | 0.6764 ± 0.0227 |
| **Chile** | Acc | 0.9513 ± 0.0079 | 0.5522 ± 0.0179 | 0.9353 ± 0.0167 | 0.9840 ± 0.0046 | 0.6777 ± 0.0095 |
| | F1 | 0.9524 ± 0.0074 | 0.6869 ± 0.0117 | 0.9391 ± 0.0150 | 0.3793 ± 0.1364 | 0.6761 ± 0.0098 |
| | AUROC | 0.9707 ± 0.0052 | 0.6838 ± 0.0229 | 0.9653 ± 0.0082 | 0.9750 ± 0.0061 | 0.7403 ± 0.0091 |
| | AUPRC | 0.9438 ± 0.0094 | 0.6725 ± 0.0160 | 0.9493 ± 0.0110 | 0.4947 ± 0.0379 | 0.7254 ± 0.0060 |
| **South Africa** | Acc | 0.9555 ± 0.0077 | 0.5440 ± 0.0170 | 0.9521 ± 0.0061 | 0.9894 ± 0.0015 | 0.6595 ± 0.0122 |
| | F1 | 0.9568 ± 0.0076 | 0.6839 ± 0.0097 | 0.9540 ± 0.0057 | 0.1508 ± 0.1496 | 0.6560 ± 0.0140 |
| | AUROC | 0.9765 ± 0.0055 | 0.7246 ± 0.0102 | 0.9759 ± 0.0043 | 0.9781 ± 0.0049 | 0.7317 ± 0.0131 |
| | AUPRC | 0.9549 ± 0.0119 | 0.7229 ± 0.0136 | 0.9624 ± 0.0059 | 0.3929 ± 0.0801 | 0.7240 ± 0.0139 |

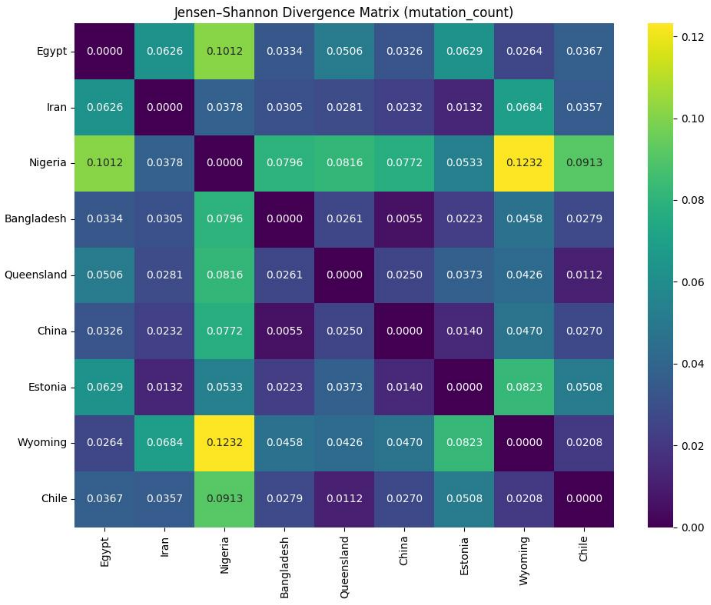


Figure 12: Jensen–Shannon divergence matrix for *mutation count* across ten regional MLGs. Higher values indicate greater dissimilarity in mutational connectivity patterns.
